# Molecular insights into early malignant transition of hepatocellular carcinoma

**DOI:** 10.1101/2025.04.14.648739

**Authors:** Zhengtao Zhang, Hong Li, Lingli Chen, Tao lu, Xinyi Shentu, Yuanhua Liu, Shuyi Ji, Zhixin Qiu, Yining Zou, Hong Wen, Jing Han, Zhengzeng Jiang, Jie Fan, Dianfan Li, Agavni Mesropian, Roser Pinyol, Josep M Llovet, Hui Dong, Yuan Ji, Lijian Hui

## Abstract

The molecular continuum of premalignant-to-malignant transition remains elusive in carcinogenesis, primarily due to the difficulty in sampling transitional lesions. Here, we present comprehensive genomic and immunological profiling of 21 very early hepatocellular carcinomas (veHCCs) arising within 17 cancer-prone DNs. Surprisingly, 82% of cancer-prone DNs harbored *TERT* alterations, suggesting a predisposing rather than causative role in malignancy transition. A substantial increase in CNA burden, rather than SNV, was noted, indicating a role of chromosome instability in malignancy transition. Additionally, veHCC-originating DNs showed immune inactivity, not falling into the prevailing paradigm that HCCs develop in a context of chronic inflammation. 43% of veHCCs showed an inflamed phenotype with relatively mild CNA burden but already exhibited immune evasion features. Two major evolutionary scenarios were thus proposed: 1) CNA-dominant progression and 2) early immune evasion of those with mild CNA burden. Collectively, our findings illustrate previously unexplored molecular paradigms in HCC initiation, highlighting the therapeutic potential of immunotherapy for early intervention.

---

Malignant transition from premalignant lesions to early cancers representing a critical stage of cancer development. For premalignant nodules, observation or survelliance would be recommended, while the diagnosis of malignancy requires immediate medical interventions(*1, 2*). Understanding this transition is essential for advancing early cancer prevention and treatment(*3*). Previous studies using evolutionarily unrelated premalignant samples and cancers only provide a statistic possibility of genetic and microenvironmental changes during the maligant transion(*4–6*). Notably, only a subset of premalignant lesions(*7*) transform to cancers, and most of them eventually regress. A large fraction of premalignant lesions thus studied would be noninformative or even misleading. Moreover, because tumor microenvironment changes dynamically, unrelated samples complicate the prediction of microenvironmental features in the malignant transition.

Apparently, only evolutionarily related premalignant and malignant samples will provide deterministic evidence. Premalignant lesions (i.e. dysplastic nodules, DNs) with early cancers developed inside, showing the nodule-in-nodule pattern, represent a snapshot of malignant transition. Nodule-in-nodule lesions would shed light on molecular events predisposing malignant transition. Additionally, these nodule-in-nodule lesions would be key to understand genetic and microenvironmental alterations that fuel the malignant transition. However, nodule-in-nodule lesions are extremely rare cases in clinical samples, because the coexisitence of DN and cancer will not last long given the increased proliferation rate of cancer cells.

Hepatocellular carcinoma (HCC), a leading cause of cancer-related mortality(*8*), exemplifies this complexity. It typically arises from DNs, which are premalignant lesions with an estimated 30% risk of transformation into HCCs(*7, 9, 10*). There have been several studies analyzed HCC initiation using evolutionarily unrelated DNs and HCCs(*4–6*). In this study, we used nodule-in-nodule lesions(*11*) as a unique opportunity to study how very early HCCs (veHCCs) develop within high-grade DNs(*12, 13*).

## Results

### Evolutionary related DNs and veHCCs from nodule-in-nodule samples

We screened 44,714 FFPE human liver specimens from 3 liver cancer centers within a 9-year interval for nodule-in-nodule samples containing DNs and veHCCs (Fig. 1A). Due to the rarity of nodule-in-nodule samples, 17 nodule-in-nodule samples from 16 individual HBV-infected patients were collected (Fig. 1A, Fig. S1, Table S1). High-grade DNs(*9, 14*) and veHCCs(*15*) were blindly and independently validated by 3 pathologists through histological evaluation of hematoxylin and eosin-stained tissue slides (Fig. 1B) and immunohistochemical (IHC) staining of GPC3, HSP70 and GS (Fig. S2A, B).

**Fig. 1.**
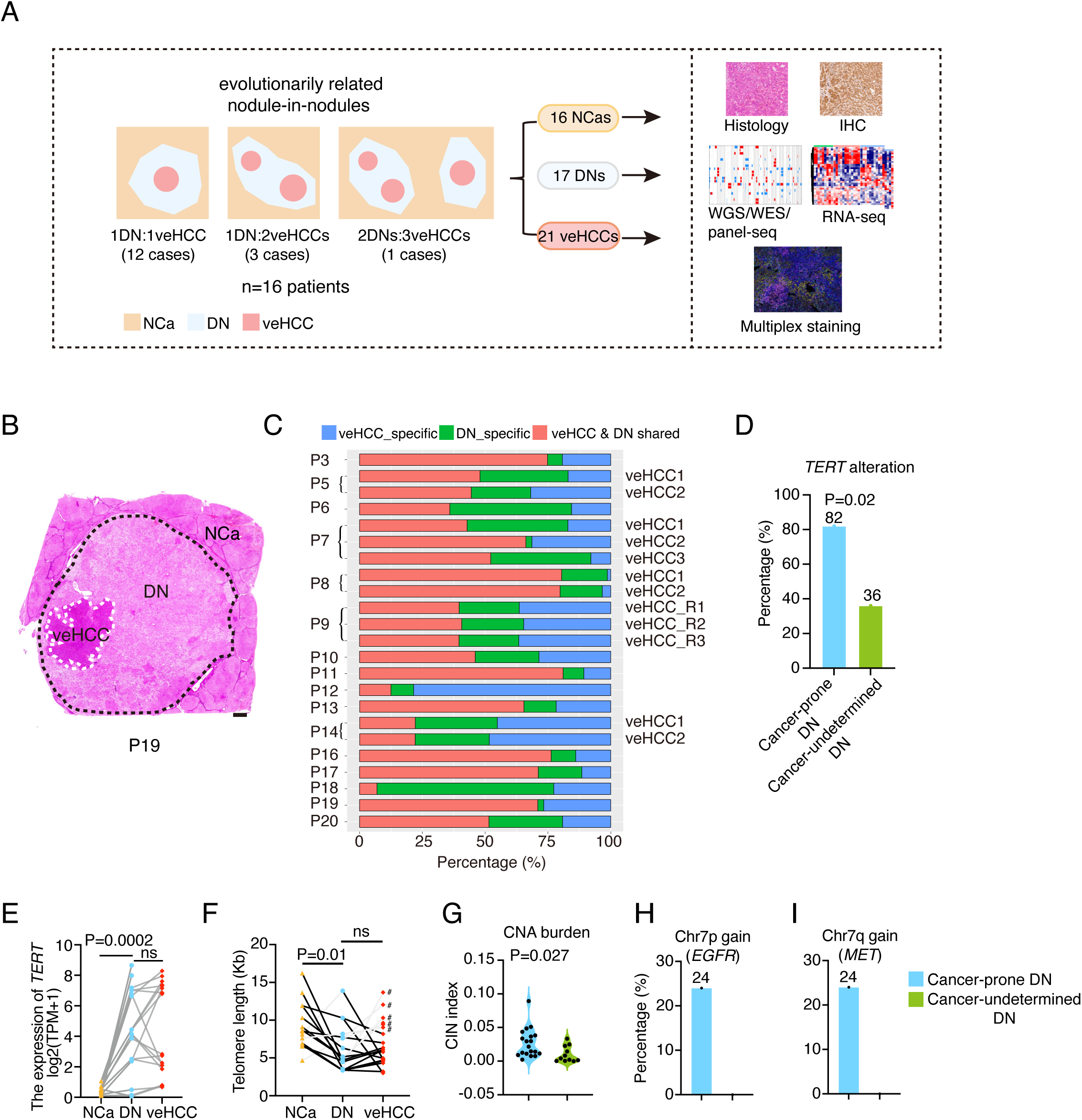
**Sample collection and characterization** (A) Schematic illustration of the experimental design for characterizing DNs and veHCCs with nodule in nodule pattern. NCa, non-cancerous tissue, DN, dysplastic nodule. (B) Whole-slide hematoxylin and eosin (H&E) images of P19. White dashed lines represent veHCC region, black dashed lines represent DN region. Scale bar, 1 mm. (C) Histogram shows the percentage of overlapped somatic single nucleotide variants (SNVs) between DN and veHCC of each case. (D) Bar plot shows the percentage of cancer-prone DNs and cancer-undetermined DNs that acquired *TERT* alterations. Significant difference is determined by Fisher’s exact test. (E) The expression of *TERT* in NCas, DNs and veHCCs based on RNA-seq data. The average expression of *TERT* of P8_DN_R1 and P8_DN_R2 were calculated and used for analysis. Those DN and veHCC samples that acquired *TERT* alterations were selected for analysis. Significant difference is determined by paired student’s t-test (two-tailed). (F) Line plots show changes in telomere length during the carcinogenesis from NCa to veHCC. Significance is determined by paired student’s t-test (two-tailed). # indicates the telomere length was extended in veHCC after malignant transition. Telomere repeat sequences (TTAGGG repeats) were obtained from WGS alignment files, and then converted to telomere length in kilobases by TelSeq. (G) Violin plot shows the chromosome instability index (CIN) of cancer-prone DNs and cancer-undetermined DNs. Significance is determined by unpaired student’s t-test (two-tailed). CNA profiles of cancer-prone DNs were inferred from WGS data, CNA profiles of cancer-undetermined DNs were inferred from WES data. (H, I) Bar plots show the percentage of cancer-prone DNs and cancer-undetermined DNs that acquired Chr7p (*EGFR*) gain (H) and Chr7q (*MET*) gain (I).

These samples were subject to WGS (about 35X) and RNA sequencing (Table S2). 17 DNs (referred as cancer-prone DNs) shared 7-85% of single nucleotide variants (SNVs) with their evolutionarily related veHCCs (21 veHCCs in total; Fig. 1C, Fig. S3A). Moreover, these DNs and veHCCs showed similar mutational signature patterns (Fig. S3B). Finally, the follow-up magnetic resonance imaging (MRI) data from patient 3 (P3) were available, which further confirmed the evolution from DN to veHCC in this case (Fig. S3C). These data collectively demonstrated their evolutionary relationship.

To confirm that SNVs could be captured from FFPE samples, we compared SNV profiles of 2 HCC samples with both freshly frozen and FFPE tissues. In line with previous reported method (*16*), 94.4% of SNVs detected in freshly frozen samples could be confirmed in FFPE tissues, showing that FFPE tissues largely retained original SNVs (Fig. S4).

### Differences between cancer-prone DN and cancer-undetermined DN

11 DNs and their paired non-cancer tissues (NCas), with no carcinoma-*in-situ* lesions, were additionally obtained from 11 HBV-infected patients. These DNs, which have undetermined cancer potential (referred as cancer-undetermined DNs), were used for comparative analysis to identify pivotal molecular events enriched in cancer-prone DNs (Table S1). Key mutations between cancer-prone and cancer-undetermined DNs were analyzed in a list of 37 liver cancer functional genes (CFGs), cataloged from 8 published HCC cohorts (Table S3)(*6, 17–23*). Remarkably, *TERT* alterations were detected in 82% of cancer-prone DNs (Fig. 1D), including HBV integration, promoter mutation and copy number gain of the *TERT* gene. By contrast, *TERT* alterations were observed in 4 of 11 cancer-undetermined DNs (36%) (Fig. 1D), which was in line with previous reported rate (*4, 5*). These data indicated a high prevalence of *TERT* alterations in cancer-prone DNs (*P*=0.02) and suggested that, in contrast to the previous conclusion(*10*), *TERT* appears to activate before the malignant transition rather than during the transition.

As previously reported(*6*), *TERT* alterations induced the upregulated expression of *TERT* in both cancer-prone DNs and related veHCCs (Fig. 1E). However, significantly shortened telomeres were still observed in 75% of cancer-prone DNs carrying *TERT* alterations (Fig. 1F). Remarkably, after malignant transition, telomere length was maintained or even shortened in 65% of veHCCs (Fig. 1F). These data showed that, different from the classic model that *TERT* alterations function to stabilize telomere length(*24*), telomere attrition was continued during the malignant transition to veHCC.

We also compared CNAs between cancer-prone and cancer-undetermined DNs. Intriguingly, significantly increased CNAs were found in cancer-prone DNs (Fig. 1G, *P*=0.027). Furthermore, there were more gains of pan-cancer oncogenes (*P*=0.02) in cancer-prone DNs (Fig. S5A, B). Notably, Chr7p and Chr7q gains, representative oncogenes including *EGFR* in Chr7p and *MET* in Chr7q, were enriched in 24% of cancer-prone DNs, but not in cancer-undetermined DNs (Fig. 1H, I). Chr8q (*MYC*) gains were also observed in 12% of cancer-prone DNs, with no occurrence in cancer-undetermined DNs (Fig. S5C). These data suggested that Chr7p/q and Chr8q gain might also contribute to the predisposition of DNs to malignant transition.

### Copy number alteration is strongly associated with malignant transition

Consistent with previous findings(*4, 6*), an increased burden of both protein-altering SNVs and CNAs was found in veHCCs when evolutionarily unrelated DNs and HCCs were compared (Fig. S6A, B). However, no significant difference was found in the numbers of total SNVs (Fig. S6C, *P*=0.57) and protein-altering SNVs (median: 52 vs 63) between DNs and paired veHCCs (Fig. 2A, *P*=0.38, Table S4), suggesting that malignant transition of cancer-prone DNs to veHCCs was not associated with SNV burden. Next, we analyzed SNV-caused CFG mutations that were relevant to the malignant transition. While new CFG mutations were detected in veHCCs, such as *ARID1A* mutations in P5_veHCC2 and P7_veHCC2, 74% of veHCCs showed no acquisition of additional SNVs in these CFGs (Fig. S6D).

**Fig. 2.**
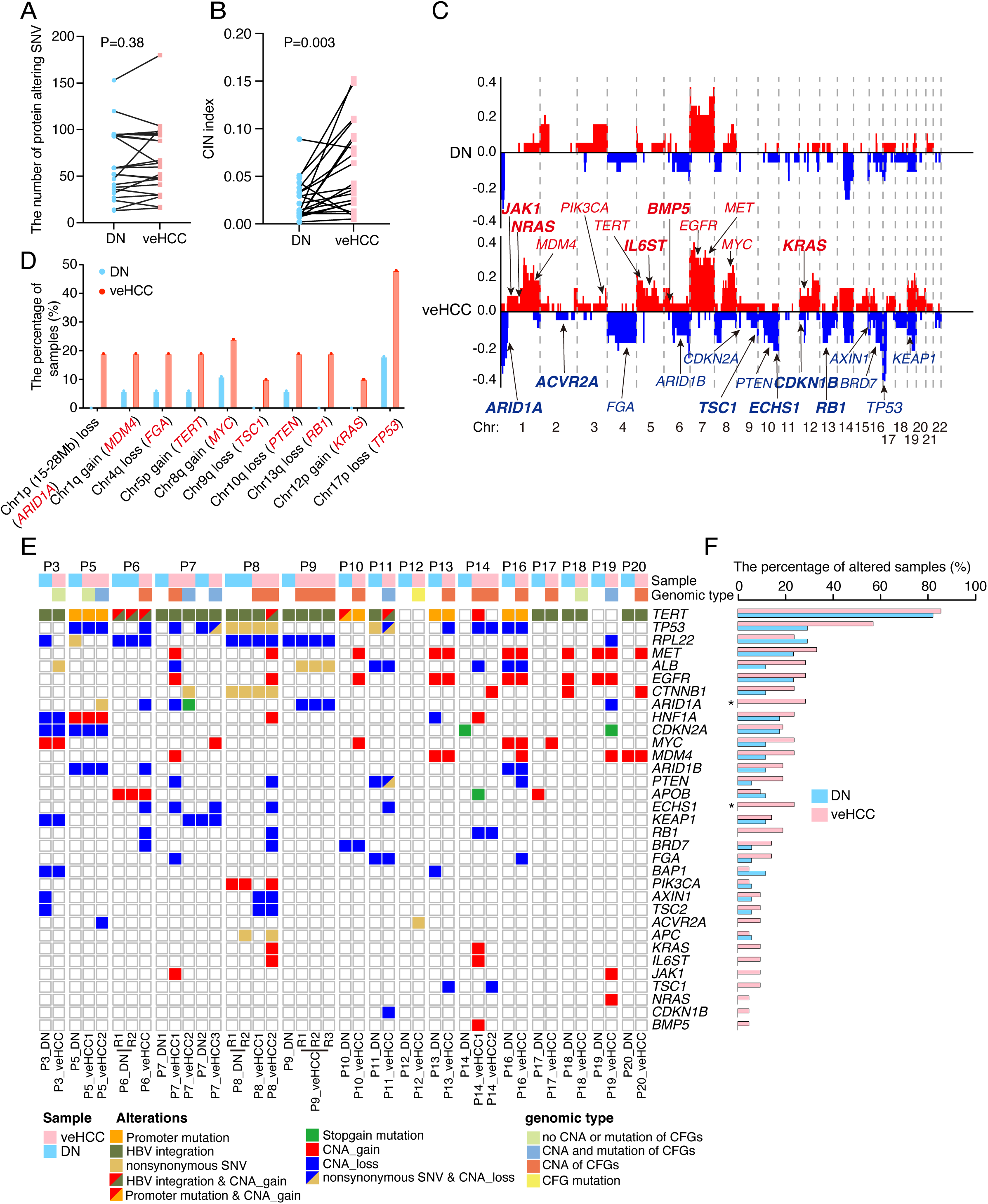
**Key genomic alterations for the malignant transition of DNs** (A) The number of protein-altering SNVs in DN and veHCC of each case detected by WGS. Significant difference is determined by paired student’s t-test (two-tailed). ns, not significant. (B) Dot plot shows the chromosome instability index (CIN) of evolutionarily related DNs and their paired veHCCs. Statistical significance is determined by paired student’s t-test (two-tailed). (C) The frequency of CNAs along the genome. The upper panel indicates the frequency of CNAs in DNs, the lower panel indicates the frequency of CNAs in veHCCs. Important liver cancer functional genes (CFGs) are labeled, with those highlighted in bold representing CFGs that are either exclusively found in veHCCs or occur with significantly greater frequency compared to those in DNs. (D) Bar plot shows the representative CNAs that exhibit a higher frequency in veHCCs compared to DNs. CFGs located within the altered genomic regions are indicated in red. (E) Heatmap shows the alteration landscape of CFGs in DNs and veHCCs. (F) Frequency of CFGs altered in DNs and veHCCs. Significant difference is determined by Fisher’s exact test. *P < 0.05.

Different from SNV burden, veHCCs exhibited significantly higher CNA burden (>1Mb) compared to their evolutionarily related DNs (Fig. 2B, Fig. S5E, F). 87% of arm-level CNAs showed increased frequency in veHCCs, including copy number losses (e.g., Chr10q (*PTEN*), Chr17p (*TP53*)) and gains (e.g., Chr1q (*MDM4*), Chr8q (*MYC*)) (Fig. 2c, d). Notably, 10 CFG-harboring CNAs were unique to veHCCs, including Chr9q (*TSC1*), Chr13q (*RB1*), Chr1p (*ARID1A*) losses and Chr12p (*KRAS*) gain (Fig. 2D).

We further expanded CNAs analysis on 803 pan-cancer oncogenes(*25*) and 1,218 pan-cancer tumor suppressor genes (TSGs)(*26*). Consistently, veHCCs harbored significantly more oncogene gains and TSG losses (Fig. S6G). Associated with increased CNA burdens, veHCCs exhibited enriched expression of chromosomal instability (CIN)-related genes (e.g., *PRC1*, *TOP2A, PTTG1*)(*27*) (Fig. S6H) and elevated DNA repair and chromosomal breakage signatures (Fig. S6I-L). These findings together highlighted the importance of CNAs in malignant transition. It is noteworthy that a veHCC-specific extended loss in Chr1p (15-28Mb) was detected in 19% of veHCCs (4/21) but absent in DNs (Fig. 2D). In their matched DNs, P6_DN and P9_DN diaplayed shorter adjacent Chr1p losses (0-15Mb) (Fig. S6M, N), suggesting progressive loss in Chr1p during malignant transition. Two HCC suppressor genes (*ARID1A, NR0B2*) in this veHCC-specific loss region showed reduced expression in veHCCs (Fig. S6O), implicating that the extended loss of Chr1p occurred during the malignant transition.

### Key genetic alterations from DNs to malignant transition

Next, integrative analysis of the SNVs, CNAs and HBV insertions was performed in 37 CFGs during the malignant transition. In total, 33 CFGs were altered in these evolutionarily related DNs and veHCCs (Fig. 2E). 22 of 33 CFGs were altered in both DNs and veHCCs (e.g., *TERT* (82% vs 86%), *TP53* (29% vs 57%), *CTNNB1* (12% vs 24%), *MYC* (12% vs 24%)) (Fig. 2F).

When SNVs and CNAs were analyzed in malignant transition-related CFGs, we found that 61% of veHCCs acquired additional CNAs in CFGs but no additional mutations in CFGs, and 22% of veHCCs accumulated both additional CNAs and mutations in CFGs. Only 1 veHCCs acquired 1 additional CFG mutation but no CNAs in CFGs (*ACVR2A*^H196Q^ in P12_veHCC). These finding again suggested that while CFG mutations contributed to the malignant transition, the acquisition of CNAs in CFGs was a major event in malignant transition (83% of veHCCs in total).

Alterations of 11 CFGs were exclusively detected in veHCCs (e.g., *ARID1A* and *ECHS1)* (Fig. 2F). *ARID1A* is a relevant TSG in HCCs(*17, 18*). However, the role of this gene in the initiation of human HCC remains unresolved. Our analysis revealed exclusive copy number loss and loss-of-function mutations of *ARID1A* in veHCCs (copy number loss in 4 veHCCs and mutations in 2 veHCCs) compared to their paired evolutionarily related DNs (Fig. 2F; *P*=0.02). *ECHS1* encodes enoyl-CoA hydratase, short chain 1, a key enzyme that functions in the mitochondrial fatty acid beta-oxidation pathway, the downregulation of which has been demonstrated to promote clear cell renal cell carcinoma growth(*28*), and *ECHS1* loss of function has been found to associated with HBV-related HCCs(*18*). Copy number loss of *ECHS1* was exclusively detected in veHCCs compared to related DN (Fig. 2E, F; 5 veHCCs vs 0 DNs), indicating its likely role in the malignant transition to veHCC.

### Immune desert phenotype in premalignant DNs

Gene expression changes were next examined during malignant transition. Differentially expressed genes (DEGs) between cancer-prone and cancer-undetermined DNs were analyzed and only 52 DEGs were identified between these 2 cohorts of DNs (Fig. S7A, *P* < 0.05). 20 of these 52 genes were significantly increased in cancer-prone DNs when compared to cancer-undetermined DNs, which were enriched for mitochondrial ATP synthesis function and small molecule metabolic process (Fig. S7B). Other 32 genes were increased in cancer-undetermined DNs, including 20 snoRNAs (Fig. S7C).

We then focused on gene expression dynamics in evolutionarily related DNs and veHCCs. Taken NCas as control, cancer-prone DNs exhibited different expression profiles from NCas analyzed by principal component analysis (PCA). Interestingly, while cancer-prone DNs showed remarkable individual specificity in genomic variations, the expression profiles of these DNs clustered to each other closely (Fig. 3A). Furthermore, cancer-prone DNs were separated from their evolutionarily related veHCCs, supporting that, despite accumulated genetic alterations, DNs have not yet developed a cancer phenotype. Notably, expression profiles of veHCCs were dispersed from one another and showed an increase in expression heterogeneity (Fig. 3A).

**Fig. 3.**
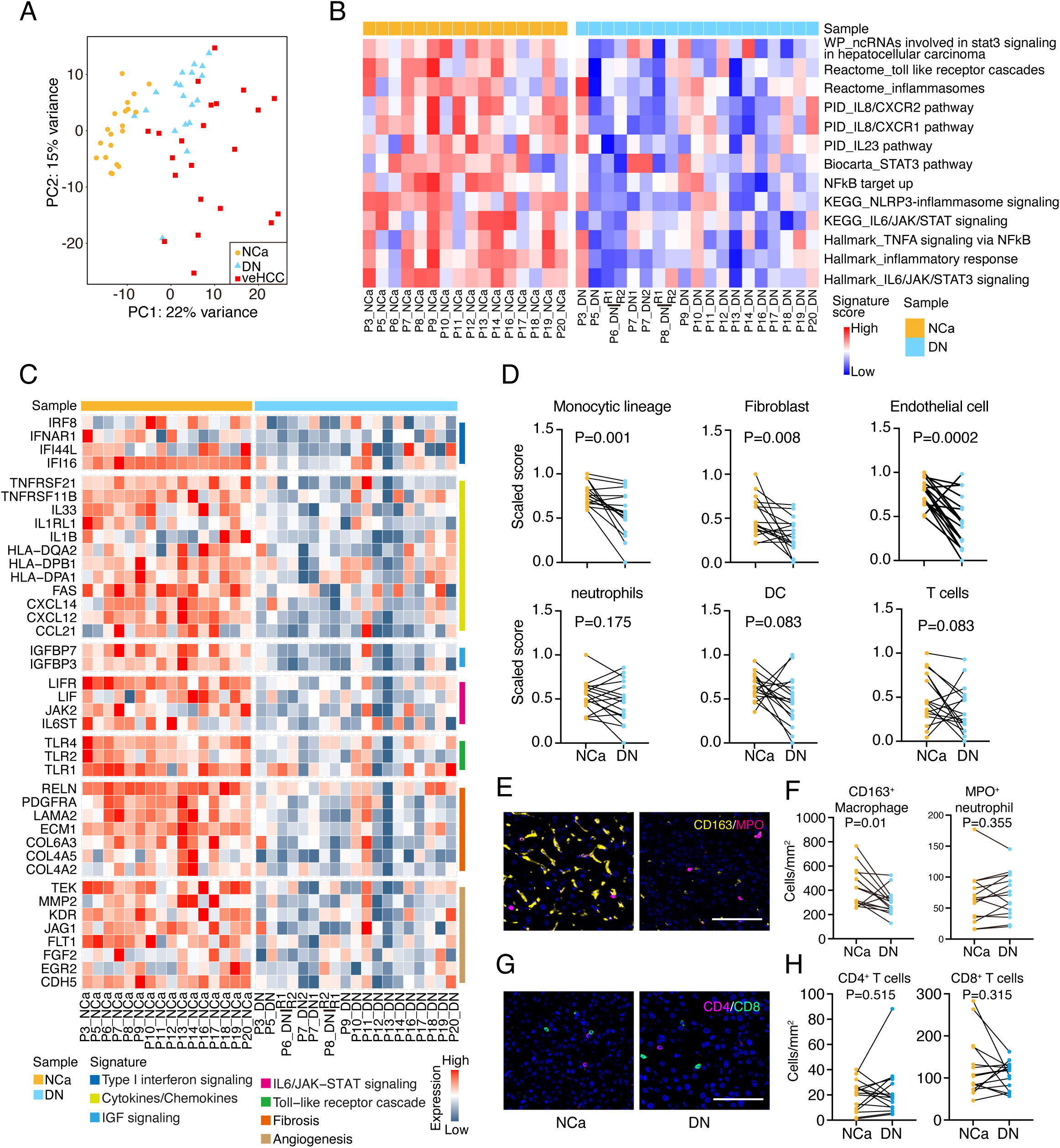
**Immune desert phenotype in premalignant DNs** (A) Principal Component Analysis (PCA) was done using top 500 variable genes among NCas, DNs and veHCCs of these evolutionarily related cases. (B) Heatmap shows the activities of inflammation associated pathways in NCas and DNs of these evolutionarily related cases. The activity of each inflammation associated pathway in each sample is determined by Gene Set Variation Analysis (GSVA). (C) Heatmap shows reduced expression of inflammation associated genes in DNs compared to respective NCas. Fold change > 2, adjust P_value < 0.05. (D) The estimation of microenvironmental cell abundance from gene-expression profiles shows the changes in immune contexture in NCas and DNs. The abundance of 6 cell types were estimated from gene expression data. DC, dendritic cell. Significant difference is determined by paired student’s t-test (two-tailed). (E) Representative immunofluorescence staining of CD163 and MPO of P11_NCa and DN. Scale bars, 100 µm. (F) The densities of CD163^+^ macrophage and MPO^+^ neutrophil was quantified as the number of cells per tissue surface area by analyzing multispectral images. Significance is determined by paired student’s t-test (two-tailed). (G) Representative immunofluorescence staining of CD4 and CD8 of P16_NCa and DN. Scale bars, 100 µm. (H) The densities of CD4^+^ and CD8^+^ T cell was quantified as the number of cells per tissue surface area by analyzing multispectral images. Significance is determined by paired student’s t-test (two-tailed).

Analyzing the DEGs further revealed three main modules that characterize the expression dynamics during the early malignancy (Fig. S7D). The largest module (777 genes), exhibiting increased expression in tumorigenesis, was associated with proliferation-related pathways (Fig. S7D, module 1). While cell cycle-related genes upregulated in DNs, the expression of the key cell cycle arrest gene *CDKN2A*, was also upregulated in DNs (Fig. S7E). In line with this finding, the number of proliferative cells was elevated in veHCCs, but not in DNs, as determined by Ki67 staining (Fig. S7F). The second module (215 genes) showed a gradually decreased expression during tumorigenesis and was enriched in pathways involved in metabolism (Fig. S7D, module 2). Among the 80 metabolism-related gene sets, 5 gene sets, including aspartate and asparagine metabolisms, were reduced in DNs (Fig. S7G), and 40 metabolism-related gene sets, including the liver metabolism of fatty acids and vitamins, were further diminished in veHCCs. Enhanced glycolysis and glucose metabolism were detected (Fig. S7H), which has been previously reported to provide favorable energy sources for cancer cells(*29*).

Interestingly, a third module (703 genes) presented a down-up expression pattern and was enriched in pathways related to epithelial-mesenchymal transition (EMT), angiogenesis and immune-related pathways, such as inflammatory response and IL2-STAT5 signaling (Fig. S7D, module 3), suggesting dynamic microenvironmental changes during the malignant transition. The activation of immune responses during HCC initiation was thus evaluated. Previous studies using mouse models demonstrated that liver cancers developed within an inflammatory microenvironment(*30–32*). However, expression of inflammation-related genes indicated that the pro-tumorigenic inflammatory signalings were reduced in cancer-prone DNs, including TNFα/NFκB, IL6/JAK/STAT3 and NLRP3-inflammasome signalings (Fig. 3B). Specifically, cancer-prone DNs showed downregulation of genes for pro-inflammatory type I interferons (*IFI44L*, *IFI16*), cytokines (*IL33*, *IL1B*), and Toll-like receptors (*TLR1*, *TLR2*, *TLR4*) (Fig. 3C, *P* < 0.05), which have been previously reported to play important roles in inflammation-related HCC development in mice(*31, 33*). Furthermore, cancer-prone DNs displayed reduced expression of genes involved in inflammation-associated fibrosis (*COL4A2*, *COL6A3*) and angiogenesis (*FGF2*, *MMP2*) (Fig. 3C, *P* < 0.05), which are known pro-tumorigenic agents in the cirrhotic and inflammatory microenvironment. Interestingly, the expression of gene signatures recapitulating inflammatory responses was also reduced in the majority of cancer-undetermined DNs (8 of 11 DNs, Fig. S7I).

Abundance of different microenvironmental cell types was estimated by deconvolution of gene expression data using MCP-counter(*34*). NCas presented high abundance scores of fibroblasts, endothelial cells and monocytic lineage cells, consistent with the cirrhotic and chronic inflammatory states of these livers(*31, 35*). However, all these types of cells were significantly reduced in cancer-prone DNs (Fig. 3D). Multiplex staining of macrophage marker CD163 further confirmed the significant reduction of macrophages in cancer-prone DNs (Fig. 3E, F). Furthermore, the infiltration of T cells was at low levels in cancer-prone DNs as determined by immunostaining of CD4 and CD8 (Fig. 3G, H). These data together indicated that cancer-prone DNs presented an immune desert phenotype, revealing an unexpected finding that HCCs originate from an immune-inactive, rather than an immune-active(*31, 33*), microenvironment.

### Immune evasion phenotype of inflamed veHCCs

Next, to characterize the immune response in veHCCs, we estimated the abundance of immune cells and calculated the activation scores of an inflamed signature of 20 core genes(*36*) in veHCCs. 43% of veHCCs exhibited both a high abundance of adaptive T cells and dendritic cells (DCs) (Fig. 4A) and high enrichment scores of the inflamed signature (Fig. 4B), which we named as inflamed veHCCs. Inflamed veHCCs displayed a significantly reduced CNA burden compared to non-inflamed veHCCs (Fig. 4C). By contrast, SNV mutation burden (Fig. S8A) and neoantigens (Fig. S8B) were comparable between these two classes of veHCCs. Integration of the transcriptional and genomic characteristics revealed two major subtypes of veHCCs: the inflamed veHCCs with low-CNA and the non-inflamed veHCCs with high-CNA.

**Fig. 4.**
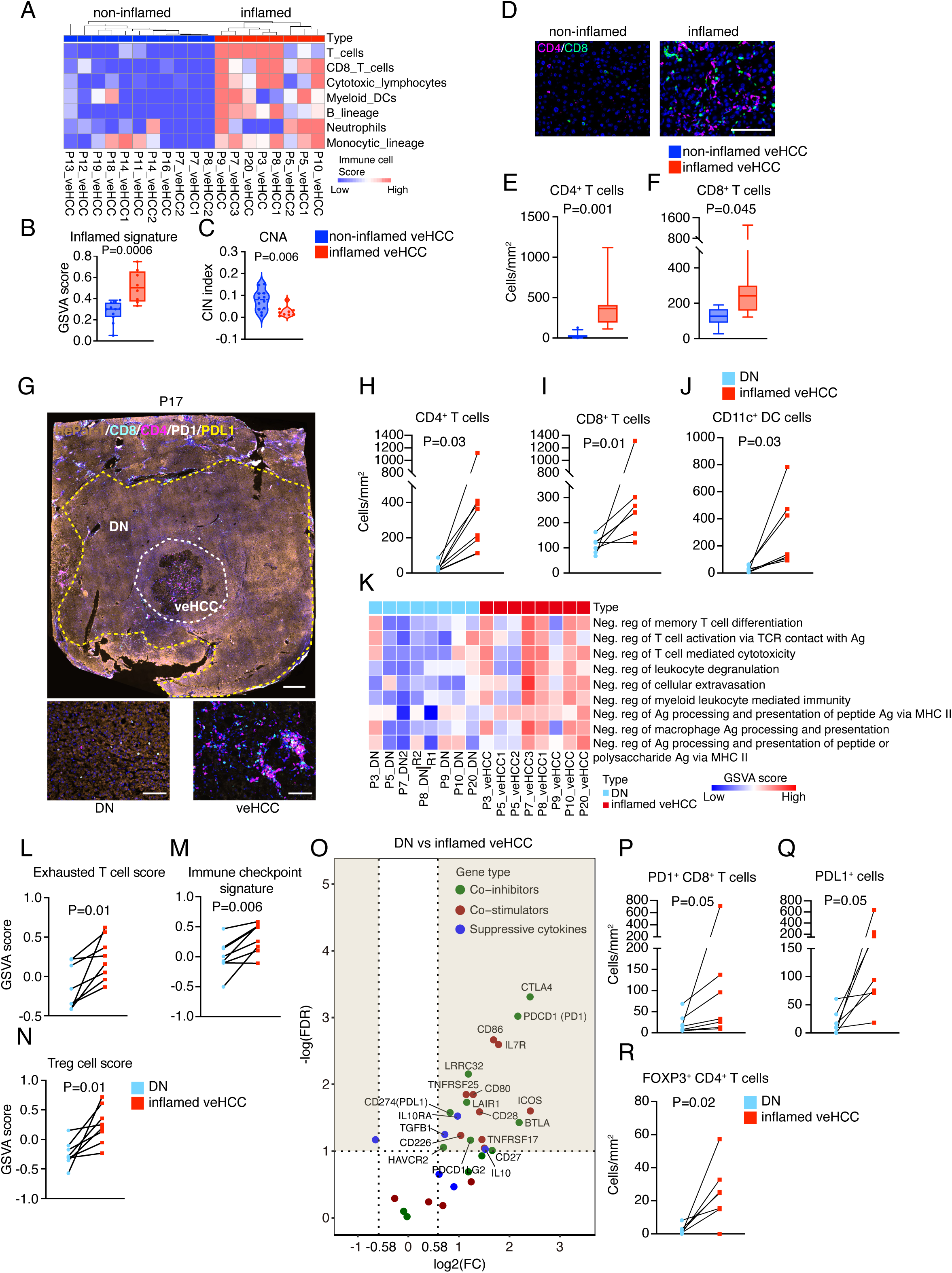
**Adaptive immune activation and evasion in a subset of veHCCs** (A) Unsupervised clustering of veHCCs based on estimated immune cell scores using the MCP-counter method. These veHCCs, which exhibited high immune cell scores, were classified as inflamed veHCCs, while the other veHCCs with low immune cell scores were classified as non-inflamed veHCCs. RNA-seq was not conducted for P6_veHCC, P9_veHCC_R2, and P17_veHCC due to insufficient material. Please be noted, P6_veHCC was classified as non-inflamed veHCC, whereas P17_veHCC was classified as inflamed veHCC based on multiplex staining data. DC, dendritic cell. (B) Box plot shows the activation scores of an inflamed signature comprising 20 genes, evaluated using GSVA. Statistical significance is determined by unpaired student’s t-test (two-tailed). (C) Violin plot shows the CNA burden in immune inflamed and non-inflamed veHCCs. Significance is determined by unpaired student’s t-test (two-tailed). (D) Representative immunofluorescence staining of CD4 and CD8 of immune inflamed veHCC (P20_veHCC) and immune non-inflamed veHCC (P16_veHCC). Scale bars, 100 µm. (E, F) Box plots show the density of CD4^+^ T cells (E) and CD8^+^ T cells (F) in immune inflamed and immune non-inflamed veHCCs. Data are presented as mean±SD. Significance is determined by unpaired student’s t-test (two-tailed). (G) Representative multiplex staining of HepPar (hepatocyte marker), PD1, PDL1, CD4 and CD8 of P17. Scale bars, 1mm (upper panel) and 100 µm (lower panel). (H-J) The densities of CD4^+^ (H), CD8^+^ (I) T and CD11c^+^ DC cells (J) were quantified as the number of cells per tissue surface area by analyzing multispectral images. Significance is determined by paired student’s t-test (two-tailed). (K) Heatmap shows the GSVA scores of gene sets associated with the negative regulation of immune responses in DNs and their matched inflamed veHCCs. The gene sets related to negative immune regulation were sourced from the Gene Ontology (GO) database. (L-M) GSVA scores of exhausted T cell signature (L), immune checkpoint signature (M) and regulatory T cell (Treg) signature (N) in DNs and paired immune-inflamed veHCCs. Significance is determined by paired student’s t-test (two-tailed). (O) Volcano plot shows the expression of co-inhibitors, co-stimulators and suppressive cytokines in DNs and paired inflamed veHCCs. FDR < 0.1 indicates significance. (P-R) Line plots show the densities of PD1^+^CD8^+^ T cell (P), PDL1^+^ cell (Q) and FOXP3^+^ CD4^+^ T cell (R) in DNs and paired immune-inflamed veHCCs. Significance is determined by paired student’s t-test (two-tailed).

Compared with the non-inflamed veHCCs, inflamed veHCCs showed increased expression of immune response-related markers, including chemokines (*CXCL12*, *CCL2*, *CCL5*), HLA class II molecules (*HLA-DRA*, *HLA-DOA*) and immune cell markers (*CD3D*, *CD4*, *CD8A*) (Fig. S8C, *P* < 0.05). Additionally, multiplex staining of CD4 and CD8 verified that the inflamed veHCCs exhibited high infiltration of CD4^+^ and CD8^+^ T cells (Fig. 4D-F). Next, we determined immune characteristics of inflamed veHCCs and compared them with their evolutionarily-related DNs. Unlike the non-inflamed veHCCs, which exhibited no enhanced immune response (Fig. S8D), inflamed veHCCs showed increased expression of adaptive immune cell markers (*CD8A*, *CD4*, *CD11c*) (Fig. S8E) and high abundance of T cells and DCs when compared to their paired DNs (Fig. S8F). Multiplex staining of CD4, CD8 and CD11c confirmed the increased infiltration of CD4^+^ T cells, CD8^+^ T cells and DCs in the inflamed veHCCs (Fig. 4G-J). Interestingly, genes involved in the bile acid pathway were downregulated in the inflamed veHCCs (Fig. S8G, H), including key enzymes *BAAT* and *CYP8B1*. Because bile acids have been shown to impede T cell responses in liver cancers using mouse models(*37*), downregulation of the bile acid pathway may be associated with the inflamed immune phenotype. It should be noted that these inflamed veHCCs were different from their inflammatory and fibrotic NCa counterparts: inflamed veHCCs showed enhanced adaptive immune responses which were missing in their paired NCas as measured by MCP-counter (Fig. S8I) and by multiplex staining (Fig. S8J). These data collectively indicated that immune responses, especially the adaptive immune responses, were enhanced in the inflamed veHCCs following malignant transition from DNs.

Surprisingly, when negative regulation of immune pathways was characterized, we found that inflamed veHCCs already upregulated the negative regulation of T cell responses and antigen processing and presentation when compared to their paired DNs (Fig. 4K). Inflamed veHCCs showed significantly high scores of immune evasion-related signatures (Table S5), including signatures for exhausted T cells (Fig. 4L), immune checkpoints (Fig. 4M), and regulatory T cells (Treg) (Fig. 4N). Signatures that recapitulate M2 macrophages and angiogenesis-associated tumor-associated macrophages (TAM) were also significantly enriched in these inflamed veHCCs (Fig. S8K). Both types of macrophages have been shown to play key roles in tumor immune evasion(*38*). Moreover, immune co-inhibitory (*CTLA4*, *PDL1*, *PD1*), co-stimulatory (*CD80*, *CD86*, *ICOS*) and suppressive molecules (*IL10*, *TGFB1*) were upregulated in these inflamed veHCCs (Fig. 4O). Consistent with the upregulation of *TGFB1*, the TGF-β signaling showed increased activation scores in these inflamed veHCCs (Fig. S8L). TGF-β has been previously reported to be associated with immune evasion(*39*) and the immune exhausted subtype of HCCs(*36*). Notably, immunostaining showed that the infiltration of PD1^+^CD8^+^ T cells, PDL1^+^ cells, FOXP3^+^CD4^+^ T cells (Fig. 4P-R) and CD163^+^ macrophages (Fig. S8M) was increased in these inflamed veHCCs. These data indicated that, on top of the inflamed phenotype, these veHCCs exhibit acquired features of immune evasion at the early stage of malignant transition.

### Integrative proposal of evolutionary scenarios of malignant transition

The data above suggested that both genomic alterations and immune microenvironmental changes occurred at early stages of HCC development. *TERT* alterations, Chr7p/7q gain and reduced immune responses were key events for the malignant transformation of cancer-prone DNs (Fig. S9). There appeared to exist two major evolutionary scenarios from DN to veHCC: the accumulation of CNAs (Scenario1) and the inflamed phenotype with immune evasion (Scenario 2) (Fig. S9). We also detected 2 outlier cases, P12_veHCC and P18_veHCC, which did not fall into these scenarios. Neither exhibited additional CNAs in CFGs nor increased immune response when compared to their evolutionarily related DNs. P12_veHCC showed one single point mutation in *ACVR2A*, whereas no additional mutations in CFGs were detected in P18_veHCC (Fig. 2E).

Phylogenetic trees were reconstructed to understand the evolutionary relationship between DN and veHCC. Somatic mutations and CNAs in CFGs were annotated on phylogenetic trees to reflect their potential role in trunk or branch evolution. CNA-mediated CFG alterations were detected in the branches of 6 cases of non-inflamed veHCCs (Fig. 5A, Fig. S10A, scenario 1). HCC development in P6 and P14 was examined as representative samples. *TERT* alteration was identified in the trunk of P6, while many CNAs were additionally observed in P6_veHCC, including Chr6q, Chr10q, Chr17p, Chr13q loss (*ARID1B*, *ECHS1*, *TP53*, *RB1*) and the extended loss of Chr1p, where *ARID1A* was located (Fig. 5A). In P14, copy number loss in Chr13q (*RB1*) and Chr17p (*TP53*) were newly acquired in the branch before the evolutionary separation of P14_veHCC1 and veHCC2. P14_veHCC1 acquired copy number gain in Chr5q (*IL6ST*), Chr6p (*BMP5*) and Chr12p (*KRAS*), and P14_veHCC2 additionally acquired copy number loss in Chr9q (*TSC1*), which may mediate the branch evolution of these 2 veHCCs (Fig. S10A).

**Fig. 5.**
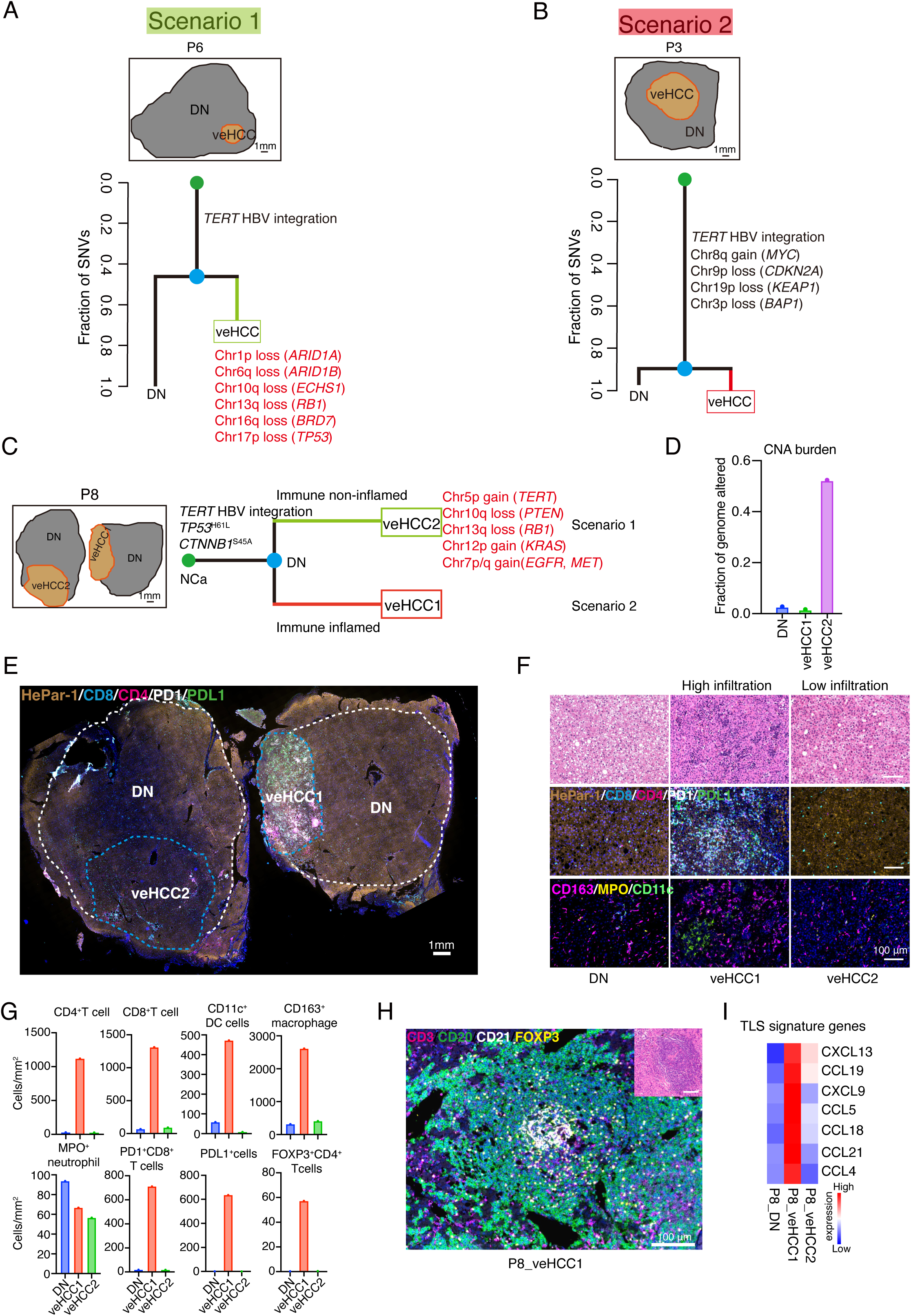
**Evolutionary scenarios of the malignant transition of DNs** (A) Spatial relationship of DN and veHCC of P6 (upper panel). Phylogenetic trees of P6 (lower panel) (Scenario 1). HBV integration in *TERT* gene and point mutations or CNAs of other CFGs were manually annotated on the trees. The trunk and branch lengths are proportional to the number of SNVs. (B) Spatial relationship of DN and veHCC of P3 (upper panel). Phylogenetic trees of P3 (lower panel) (Scenario 2). (C) Schematic illustration of the 2 evolutionary scenarios of the malignant transition from P8_DN to P8_veHCC1 and P8_veHCC2. HBV integration in *TERT* gene and point mutations or CNAs of other CFGs were manually annotated on the trees. (D) Bar plot shows the CNA burden in the DN and paired veHCCs of P8. (E) Multiplex staining of HepPar, PD1, PDL1, CD4 and CD8 of P8. Scale bar, 1mm. (F) Representative H&E staining (upper panel), multiplex staining of HepPar, PD1, PDL1, CD4, CD8 (middle panel) and multiplex staining of CD163, MPO, CD11c (bottom panel) of P8. Scale bars, 100 µm. (G) Bar plots show the densities of CD4^+^ and CD8^+^ T cells, CD11c^+^ DC cells, CD163^+^ macrophage, MPO^+^ neutrophil, PD1^+^ CD8^+^ T cells, PDL1^+^ cells and FOXP3^+^ CD4^+^ T cells in DNs and veHCCs of P8. (H) Multiplex staining of CD3, CD20, CD21 and FOXP3 of tertiary lymphoid structure (TLS) in P8_veHCC1. H&E staining shows the histology of TLS. Scale bars, 100 µm. (I) Heatmap shows the high expression of TLS-related signature genes in P8_veHCC1.

Phylogenetic trees of 9 inflamed veHCCs were analyzed as scenario 2. 6 of these 9 inflamed veHCCs (P5_veHCC2, P7_veHCC3, P9_veHCC, P10_veHCC, P17_veHCC, and P20_veHCC) showed copy number losses or gains in CFGs compared to their paired DNs (Fig. S10B, C, Scenario 2); however, CNA alterations in these veHCCs were low compared with those in scenario 1. Remarkably, the other 3 inflamed veHCCs (P3_veHCC, P5_veHCC1, P8_veHCC1) exhibited nearly identical CNAs compared to their paired DNs. Moreover, they did not acquire additional mutations in the 37 CFGs (Fig. 5B, Fig. S10B, C). We further analyzed mutations in previously reported 803 oncogenes(*25*) and 1,218 TSGs(*26*). Remarkably, no additional point mutations in these genes were identified in P8_veHCC1. In P5_veHCC1, nonsynonymous mutations were detected in *EEF1A1*, *NDST4* and *CTNNA2*, and in P3_veHCC, a point mutation in *UFL1* was observed. Interestingly, *NDST4* and *CTNNA2* showed very low expression levels in both P5_DN and their paired veHCCs, and the roles of *EEF1A1* and *UFL1* in liver carcinogenesis remain unclear. These findings suggested that major cancer genes appeared to be missing in these veHCCs, leaving a possible contribution of immune evasion to malignant transition of the 3 veHCCs.

We observed 2 cases (P7 and P8), where spatially separated veHCCs were developed from different evolutionary scenarios within the same patients (Fig. S10C). P7 represented a typical clinical case in which multifocal HCCs were independently initiated from spatially separated DNs (Fig. S11). Intriguingly, in P8, 2 spatially separated veHCCs representing 2 evolutionarily different scenarios were developed from the same DN (Fig. 5C). P8_veHCC2 showed a large number of CNAs, covering 12 chromosomes, which resulted in gain of oncogenes (*EGFR*, *MET, KRAS*) and loss of tumor suppressors (*PTEN*, *RB1*) (Fig. 5C, D), suggesting that the high burden of CNAs could potentially explain the malignant transition to P8_veHCC2 (Scenario 1). By contrast, P8_veHCC1 presented almost identical CNAs compared to P8_DN (Fig. 5D), and no additional mutations in CFGs were detected in P8_veHCC1, even after tripling the sequencing depth of P8_veHCC1 compared to the paired DN (Table S2). However, different from P8_veHCC2, P8_veHCC1 showed an inflamed phenotype compared to P8_DN (Fig. 5E-G). Histological assessment, multiplex staining (CD3-T cells, CD20-B cells, CD21-follicular dendritic cells and FOXP3-Treg cells) and signature expression analysis suggested the presence of tertiary lymphoid structure (TLS) in P8_veHCC1 (Fig. 5H, I). TLS are key players in the response to checkpoint inhibitor-based immunotherapies(*40–42*) and have been shown to promote HCC initiation(*43*). Furthermore, the increased infiltration of PD1^+^CD8^+^ T, PDL1^+^ and FOXP3^+^CD4^+^ T cells (Fig. 5G) might mediate immune evasion and facilitate the malignant transition of P8_DN to P8_ veHCC1 (Scenario 2).

## DISCUSSION

Unlike previous publications(*4, 5*), our study focused on evolutionarily related DNs and veHCCs, offering unique insights into the spatiotemporal genomic, transcriptomic, and immune microenvironmental changes during malignant transition to veHCCs. Our findings provide direct evidence into the molecular drivers and evolution scenarios of HCC malignant transition, which challenge the existing paradigms of cancer development in several aspects (see below). While our cohort size remains limited, our analysis of the CFGs and CNAs indicates that our samples covered most recurrent CFGs (e.g., *TERT*, *TP53*, *CTNNB1*) and CNAs (e.g., Chr1q, Chr8q gain and Chr10q, Chr17p loss) observed in malignant HCCs, strongly suggesting their representative. Furthermore, several biologically striking findings emerged. Most notably, *TERT* alterations and diminished immune responses were detected in over 80% of DNs, suggesting these lesions acquire oncogenic “priming” before histological malignancy. Furthermore, genomic and immune profiling revealed distinct evolutionary scenarios during early hepatocarcinogenesis: chromosomal instability manifested as increased CNA burden was observed in over 80% of veHCCs, while approximately 50% of veHCCs concurrently exhibited an immunosuppressive tumor microenvironment. These parallel findings strongly implicate chromosomal instability and immune microenvironment remodeling as dual drivers of the premalignant-to-malignant transition. It should be acknowledged that our sampling strategy focused on HCCs arising through the DN-HCC sequence, potentially excluding de novo carcinogenesis pathways not involving detectable DN precursors.

Telomerase activation caused by *TERT* alterations is a hallmark of HCC, present in over 90% of cases(*10, 24*). Interestingly, *TERT* alterations were previously observed in 20% of undetermined DNs, which included those either progressing to HCCs or eventually regressing, raising the possibility that *TERT* alterations might be a driver of malignant transition(*4, 5*). However, *TERT* alterations were detected in 82% of cancer-prone DNs, indicating that *TERT* alterations may not be direct drivers of malignant transition but instead potentially function as gatekeepers. All nodule-in-nodule lesions in this study occurred in the background of cirrhosis. Recent studies have identified *TP53* and *ARID1A* mutations but no *TERT* alterations in cirrhotic liver tissues(*44, 45*). Taking this into account, results from the analysis of DNs(*44*) suggest that mutations in other cancer-related genes may not be as essential as *TERT* for the genesis of cancer-prone DNs. Additionally, our data suggests the possibility for re-evaluating the functions of *TERT* alterations in the malignant transition of other cancer types, such as melanoma, glioma and bladder cancer(*46*).

Human liver cancers are characterized by numerous genomic alterations, including SNVs and CNAs(*17, 18*). However, the sequential order of these alterations and their role in malignant transition remain largely unknown. We found that the SNV burden does not increase, and acquisition of additional mutations in cancer-related genes are less common compared to CNAs during the malignant transition. This shifts the focus away from the conventional view that driver gene mutations are the major instigators of cancer development(*4, 5, 10, 13*). Instead, we show that CNAs frequently accumulate during the malignant transition. Copy number gains in oncogenes and losses in TSGs play a pivotal role in driving the transition from DN to veHCC. We show that pathways related to chromosomal instability, chromosome breakage, and DNA repair were remarkably enhanced in veHCCs, suggesting that genomic instability may underlie these CNA-driven changes. Notably, while *TERT* expression persisted throughout the transition, telomere attrition was not completely prevented during the transition from DN to HCC. This may be also correlated with chromosome instability.

It is unclear how the immune microenvironment evolved and participated in the transition from premalignancy to malignancy. Surprisingly, we found that DNs are dominated by an immune desert phenotype where immune responses were inactivated. Previously, inflammation has been demonstrated to enhance liver carcinogenesis by providing excessive cytokines and growth factors in microenvironment(*30–32*). The fact that veHCCs evolve from immune inactive DNs suggests that there is a need for reevaluating the role of inflammation in liver carcinogenesis. Notably, when immune states of veHCCs were analyzed in this study, a subset of veHCCs showed an immune-inflamed phenotype (“inflamed veHCCs”). Interestingly, these veHCCs already acquired immune evasion-related features, suggesting the concurrence of an inflamed status and immune evasion at the very early stage of HCC initiation. These data pointed out the possibility that an immune evasion state might contribute to the malignant transition.

Strikingly, we observed two major evolutionary scenarios from DN to veHCC: one with high CNA burden (Scenario 1), and another with an immune inflamed phenotype displaying features of immune evasion (Scenario 2). Among the cases in Scenario 2, P3_veHCC, P5_veHCC1 and P8_veHCC1 exhibited nearly identical CNAs compared to their paired DNs. Furthermore, no SNVs in known HCC driving genes or pan-cancer genes were acquired in P8_veHCC1. P3_veHCC only presented one additional SNV in the *UFL1* gene. A recent structural study(*47*) revealed that Ser205 lies at the interface between UFL1 and the 60S ribosome. However, given that the interface surface is broad, it is hard to tell whether *UFL1*^S205A^ mutation may impact on ribosomal binding. In P5_veHCC1, three new SNVs were identified in the *EEF1A1*, *NDST4*, and *CTNNA2*. *NDST4* and *CTNNA2* were expressed at very low mRNA levels and may not involve in HCC formation. AlphaFold2(*48*) predicted that residue Asn284 is located on the protein surface, away from the GTP-binding site based on its structural similarity to GTP-bound GTPases(*49*). Thus, it is unlikely that the *EEF1A1*^N284K^ mutation directly affects the GTPase activity of eEF1A1. Although it is premature to draw definitive conclusions, these findings collectively suggest a testable hypothesis: in some cases, the malignant phenotype may arise due to factors other than genetic alterations(*50*). Supporting this hypothesis, one recent study has demonstrated that transient loss of Polycomb components is sufficient to initiate tumorigenesis in *Drosophila* without the involvement of cancer driver mutations(*51*). In line with this direction, it is exciting to ask whether some cancer cells, even after acquiring mutations, could be converted back into a non-malignant states.

Our findings provide several insights that could be valuable in clinical practice. Firstly, the detection of *TERT* alterations as potential markers for cancer-prone DNs offers an opportunity for early stratification and non-invasive monitoring through circulating DNA analysis. Previously, *TERT* promoter mutations have been detected in the circulating DNA of HCC patients(*52*), however further efforts are needed to translate these findings into clinical applications. Moreover, the high prevalence of TERT overexpression might serve as a possible target for immunotherapy for eliminating cancer-prone DNs and veHCCs(*53*). Secondly, exploring CNAs and their associated genomic instability pathways might potentially open new avenues for therapeutic intervention during the transition to malignancy. Synthetic lethal targets have been identified to exploit the vulnerabilities of cancers with specific chromosomal alterations, including the loss of chromosome 1p(*54*), 18q or 16q(*55*), deletions at chromosome 9p21.3(*56, 57*). Thirdly, the high abundance of CD8^+^ T cells, the formation of TLS, an overexpression of inflamed signature genes, along with the upregulation of PD-1 and PD-L1, suggests that immune checkpoint inhibitors could be beneficial for patients with inflamed veHCCs. Additionally, the increased infiltration of Treg cells and enhanced TGF-β and angiogenesis signaling in inflamed veHCCs indicate the potential for combination therapies to improve efficacy or overcome resistance to anti-PD-1 or anti-PD-L1 treatments.

## Acknowledgments

We thank Chenqi Xu, Jinqiu Zhou (Center for Excellence in Molecular Cell Science, CAS), Bing Li (Shanghai Institute of Immunology, Shanghai Jiaotong University), Pengyuan Yang (Institute of Biophysics, CAS), Lin Liu (Nankai University) and Denggui Li (Suzhou Institute of Systems Medicine) for helpful discussions and suggestions. We thank all the patients for their consent and participation in this study.

## Funding

This study is supported by the National Natural Science Foundation of China (NSFC) (92168202, 91942313, 32000578, T2122018, 32470707), Shanghai Municipal Science and Technology Major Project, the “Strategic Priority Research Program” of the Chinese Academy of Sciences (XDA16020201), the National Key Research and Development Project (2019YFA0802001). JML was supported by grants from European Commission (Horizon Europe-Mission Cancer, THRIVE, Ref. 101136622), the NIH (R01-CA273932-01, R01DK56621 and R01DK128289); Samuel Waxman Cancer Research Foundation; the Spanish National Health Institute (MICINN, PID2022-139365OB-I00, funded by MICIU/AEI/10.13039/501100011033 and FEDER); Cancer Research UK (CRUK), Fondazione AIRC per la Ricerca sul Cancro and Fundación Científica de la Asociación Española Contra el Cáncer (FAECC) (Accelerator Award, HUNTER, Ref. C9380/A26813); the “la Caixa” Banking Foundation; Fundación Científica de la Asociación Española Contra el Cáncer (FAECC; Proyectos Generales, Ref. PRYGN223117LLOV; and Reto AECC 70% Supervivencia: Ref. RETOS245779LLOV) and the Generalitat de Catalunya/AGAUR (2021 SGR 01347). R. Pinyol was supported by the Fundació de Recerca Clínic Barcelona - IDIBAPS and by a grant from the Spanish National Health Institute (MICINN, PID2022-139365OB-I00, funded by MICIU/AEI/10.13039/501100011033 and FEDER).

### Author contributions

Conceptualization, Zhengtao Zhang, Lijian Hui, Hong Li, Yuan Ji; Methodology, Zhengtao Zhang, Hong Li, Lingli Chen, Shuyi Ji, Dianfan Li; Software, Hong Li, Xinyi Shentu, Yuanhua Liu; Experiments, Zhengtao Zhang, Zhengzeng Jiang; Sample and Histology, Yuan Ji, Lingli Chen, Hui Dong, Tao Lu, Hong Wen, Jie Fan, Jing Han, Yining Zou; Writing, Zhengtao Zhang, Lijian Hui, Hong Li, Agavni Mesropian, Roser Pinyol, Josep M Llovet, Yuan Ji, Zhixin Qiu; Supervision, Lijian Hui, Yuan Ji, Hong Li.

### Data availability

The raw sequence data reported in this paper have been deposited in the Genome Sequence Archive (Genomics, Proteomics & Bioinformatics 2021) in National Genomics Data Center (Nucleic Acids Res 2022), China National Center for Bioinformation / Beijing Institute of Genomics, Chinese Academy of Sciences (GSA-Human: HRA008950) that are publicly accessible at https://ngdc.cncb.ac.cn/gsa-human.

**Supplementary materials**

## Materials and Methods

Fig. S1 to S11

Table S1 to S5

