## Supplementary Figures for "Molecular insights into early malignant transition of hepatocellular carcinoma"

**The PDF file includes:**

Materials and Methods
Supplementary table legends
Supplementary figure legends
Figure S1-S11

### Materials and Methods

#### Patient cohort

These samples were collected from the department of pathology, Zhongshan Hospital, Fudan University; department of pathology, Eastern Hepatobiliary Surgery Hospital, Naval Medical University and department of pathology, Huashan Hospital, Fudan University. This study was approved by the Ethics Committee Board at Zhongshan Hospital, Eastern Hepatobiliary Surgery Hospital and Huashan Hospital.

More than 44,714 formalin-fixed paraffin-embedded (FFPE) liver specimens, derived from surgically resection of liver cancers within a 9-year interval (2014-2022), were screened to identify coexistent high grade dysplastic nodules (DNs) and very early hepatocellular carcinomas (veHCCs) with nodule-in-nodule pattern. To diagnose high grade DN and veHCC, pathological reviewing was independently done by three expert pathologists for each case. The following histological features were quoted: mean of nodule sizes; steatosis; unpaired arteries; portal tract; stromal invasion; hepatic plate thick; inflammatory infiltrate; ductular reaction; cytological atypias; pseudoglandular formation; cholestasis; clear cell changes; ballooning; hyaline inclusions; mallory bodies; and large and small cell changes. Diagnosis was based on WHO defined histological criteria for hepatocellular neoplasia and followed the published method (5, 9). In case of diagnostic difficulties, the pathologists have used the results of immunostaining of GPC3, HSP70 and GS. In this situation, positivity of at least two markers was considered sufficient to diagnose early HCC. Furthermore, due to very early stage of these HCCs, we classified these nodule-in-nodule HCCs as veHCCs

(15). In total, 17 evolutionarily related nodule-in-nodule lesions from 16 patients (1 lesions from each patient of P3, P5, P6 and P8-P14, P16-P20, and 2 spatially separated lesions from P7) were collected. Please be noted, although P8 has two spatially separate samples, both samples are from the same DN. These nodule-in-nodule high-grade DNs and their paired veHCCs were taken by microdissection and then subjected to whole genome sequencing (WGS, ~35X) and RNA sequencing (RNA-seq). 4 cases (P1, P2, P4 and P15), although spatially overlapping, exhibited less than 1% overlap in single nucleotide variants (SNVs) and no shared copy number alterations (CNAs) between DNs and veHCCs, indicating that they originated independently. Additionally, 7 individual high-grade DNs were also collected from 7 patients. We grouped these 11 DNs and classified them as cancer-undetermined DNs. 15 individual eHCCs were collected from 15 patients. These evolutionarily unrelated DNs and eHCCs were subjected to whole exome sequencing (WES, ~230X), target sequencing (e.g., TERT promoter, CTNNB1, TP53 and ARID1A) (~1880X), and RNA-seq. CNAs and protein-altering SNVs were identified by analyzing the WES data, TERT promoter alterations (promoter mutation and HBV integration) were further identified by analyzing the panel sequencing data. All patients had hepatitis B virus (HBV) infections and developed cirrhosis aged from 32 to 72. Table S1 summarized the detailed information of tissue coding and clinicopathological data.

**Preparation of tissues for sequencing**

Following the acquisition of formalin-fixed paraffin-embedded (FFPE) blocks of the lesions from the tissue bank, serial sections were prepared at a thickness of 5 μm from each lesion for H&E staining and immunostaining procedures. In the case of nodule-in-nodule lesions, a thorough pathological review was conducted on the sections of each lesion to delineate the areas corresponding to DNs and veHCCs. For individual DN or eHCC lesions, a similar pathological review was performed to accurately identify the respective areas of DN or eHCC. Subsequently, DN and eHCC tissues were meticulously harvested through microdissection. Additionally, non-cancerous tissue (NCa) for each case were also obtained via microdissection to enrich parenchymal cells instead of removing the fibrous bands.

The harvested tissues were transferred to 1.5 mL microcentrifuge tube and deparaffinization solution was used to remove paraffin. The Maxwell® 16 LEV RNA FFPE Purification Kit (Promega) was used to extract FFPE RNA according protocol instructions. RNA integrity was determined by 2100/2200 Bioanalyzer (Agilent) with DV200 (Percentage of RNA fragments > 200 nt fragment distribution value) and quantified using the NanoDrop (Thermo Fisher Scientific). For FFPE DNA extraction, The Maxwell® 16 FFPE Plus LEV DNA Purification Kit (Promega) was used according to protocol instructions. The integrity and concentration of the total DNA was determined by agarose electrophoresis and Qubit 3.0 fluorometer dsDNA HS Assay (Thermo Fisher Scientific).

**RNA Sequencing and analysis**

RNA purification, reverse transcription, library construction and sequencing were performed according to the manufacturer's protocol (Illumina). The captured coding regions of the transcriptome from total RNA were prepared using TruSeq® RNA Exome

Library preparation Kit. For FFPE sample, RNA input for library construction was determined by the quality of RNA. Generally, 20 ng RNA was recommended for high quality RNA and 20-40 ng RNA for medium quality RNA. Then the cDNA was generated from the input RNA fragments using random priming during first and second strand synthesis and sequencing adapters were ligated to the resulting double-stranded cDNA fragments. The coding regions of the transcriptome were then captured from this library using sequence-specific probes to create the final library. After library constructed, Qubit 3.0 fluorometer dsDNA HS Assay (Thermo Fisher Scientific) was used to quantify the concentration of the resulting sequencing libraries, while the size distribution was analyzed using Agilent Bioanalyzer 2100 (Agilent). Paired-end 150 bp sequencing was performed on NovaSeq 6000 S4 sequencer following the protocol instructions (Illumina) in Mingma Technologies at Shanghai, China. RNA-seq was not performed for P6_veHCC and P17_veHCC due to insufficient tissue availability.

Raw reads were first trimmed to remove the low-quality bases and adapter sequences. High-quality reads were mapped to human GRCh38 genome by TopHat2 algorithm (58). Counts of individual transcripts were quantified using FeatureCounts. Differential analysis across groups were performed using DESeq2 with the count data as the input, and the batch and individual information were involved in the design to compensate the batch effect and individual difference. Genes with the absolute change >=2 and adjust P value less than 0.05 were chosen as significant. For PCA analysis of all evolutionarily related samples, batch effect of the counts was removed using the function ComBat_seq in R package sva and then transformed using vst from DESeq2.

PlotPCA in DESeq2 was used to extract the principal components with the top 500 variable genes.

**Pathway activity and enrichment analysis**

Pathway activities were estimated by GSVA (59). Cancer hallmark pathway-related (Fig. S7D), DNA repair-related (Fig. S6J-L), Reactome metabolism-related (Fig. S6G, H), inflammation-related gene sets (Fig. 3B, Fig. S7I), and GO negative immune regulation-related (Fig. 4K) were downloaded from MSigDB website for analysis. Immune evasion-related signatures, including signatures for exhausted T cells (Fig. 4L), immune checkpoint (Fig. 4M), and Treg (Fig. 4N) and TGF-β signature (Fig. S8L) were collected from published studies (See Table S5 for detailed information). A list of 70 chromosome instability associated signature genes (27) (Fig. S6H) and chromosome breakage-related gene set (Fig. S6I) from Human Phenotype Ontology Data base were analyzed by pre-ranked Gene Set Enrichment Analysis.

**Deconvolution of immune cell composition based on gene expression**

Relative proportion (abundance) of immune cell was estimated by Microenvironment Cell Populations (MCP)-count (34) from gene expression matrix. MCP count produces for each sample an abundance score for CD3+ T cells, CD8+ T cells, cytotoxic lymphocytes, NK cells, B lymphocytes, cells originating from monocytes (monocytic lineage), myeloid dendritic cells, neutrophils, as well as endothelial cells and fibroblasts.

**Whole genome sequencing and analysis**

The DNA was sheared to an average target size of 350 bp using Covaris S220 Sonicator (Covaris). Fragmented DNA was purified using Sample Purification Beads

(Illumina). Adapter-ligated libraries were prepared with the TruSeq Nano DNA Sample Prep Kits (Illumina) according to protocol instructions (Illumina).

DNA concentration of the enriched sequencing libraries was measured by the Qubit 3.0 fluorometer dsDNA HS Assay (Thermo Fisher Scientific). Size distribution of the resulting sequencing libraries was analyzed using Agilent Bioanalyzer 4200 (Agilent). Paired-end sequencing is performed on the Illumina HiSeq X TEN or NovaSeq 6000 system with 2x150 bp paired-end reads following protocol instructions (Illumina).

Raw reads of WGS were processed by an in-house script to remove low-quality reads and trim adapters (60). The remained reads were mapped to human genome (hg19) by BWA algorithm (61). Duplicate reads were removed by Picard tools. SNVs and indels were called and recalibrated by Genome Analysis Toolkit (GATK) (62). Variants were hard-filtered using the parameters recommend by GATK Best Practices. Similarity between any paired samples were calculated based on the Jaccard similarity coefficient between their germline variants. Hierarchical clustering analysis were performed among all samples to check the source of samples.

Somatic single nucleotide variations (SNVs) were called by muTect (63) and varScan2(64). Initial mutation calls were filtered by the recommend criteria of each algorithm. To obtain more confident somatic mutations, only those mutations identified by two algorithms or occurred in more than one sample were kept for downstream analysis. ANNOVAR (65) was used to map mutations to proteins to estimate the potential functional effects. Please be noted, to avoid missing true mutations in protein-altering mutations, mutations identified by either of the two tools were manually checked. 37 CFGs in this study were collected based on significantly mutated HCC genes identified across 8 genome sequencing studies (6, 17-23). This includes 19 genes that were reported in more than 2 studies: TERT, TP53, CTNNB1, AXIN1, ALB, BAP1, KEAP1, NFE2L2, RB1, PIK3CA, RPS6KA3, KRAS, IL6ST, CDKN2A, ARID1A, ARID2, ACVR2A, APOB and NRAS. Additionally, 12 genes frequently mutated in HBV-related HCCs (18) were included: TSC1, TSC2, JAK1, BRD7, FGA, PTEN, HNF1A, PRDM11, CDKN1B, BMP5, RPL22 and ECHS1. We also included 6 additional genes that play important roles in HCC development: MET, EGFR, APC, MDM4, MYC and ARID1B. Table S3 summarizes the list of CFGs. CFG mutations identified by either of the two tools were manually checked and potential false mutations were removed: germline mutations that were frequent in population, especially in Asians; mutations occurred in reads containing more than 3 different base changes; mutations with obvious strand bias; cluster-mutations occurring in a certain region; mutations covered by less than 3 different reads.

Genome sequence of HBV was downloaded from NCBI (NC_003977.2). HBV integration breakpoints were identified and annotated by Virus-Clip (66). Sequencing reads were aligned to the HBV genome. Then soft-clipped reads were extracted to identify human and HBV integration breakpoints.

Somatic copy number alterations (CNAs) were identified by BIC-Seq (67) software. BIC-Seq returned log2 ratios between the observed and expected number of reads in the segments. Segments with log2 ratio >0.2 or <−0.2 were regarded as Somatic CNAs. SNVs with insufficiency depth (<15) were removed. The remained SNVs were passed to Pairtree (68, 69) and phylogenetic trees were built by Pairtree from multiple samples of each patient.

Telomere repeat sequences (TTAGGG repeats) were obtained from WGS alignment files, and then converted to telomere length in kilobases by TelSeq (70).

**Whole exome sequencing and Panel sequencing**

Whole exome sequencing (WES) libraries were prepared and captured using the SureSelectXT Human All Exon V8 kit (35.1 Mb, Agilent Technologies) following manufacturer’s instructions. The DNA library was sequenced with Illumina NovaSeq 6000 system. After removing low-quality reads and trim adapters (60), the remained reads were mapped to the human genome (hg19) by BWA algorithm(61). Duplicate reads were removed by Picard tools and recalibrated by GATK (62). SNVs were called by muTect (63) and varScan2 (64). ANNOVAR (65) was used to map mutations to proteins to estimate the potential functional effects. After these analysis procedures, the protein-altering SNVs identified by either of the two tools were manually checked and a downstream filter comprised of the following criteria was used to obtain high quality SNVs: (1) Coverage > 8X; (2) Variant Allele Frequency (VAF) >= 5% and at least 4 variant supporting reads in the DN or eHCC samples, and VAF < 1% and no more than 2 variant supporting reads in non-cancerous tissues; (3) mutations identified in the snp138 database were removed. CNAs were identified by comparing eHCC or DN samples with their paired non-cancerous tissues using CNVkit (71).

A 65-gene panel was designed, encompassing important liver cancer functional genes, e.g., TP53, CTNNB1, ARID1A, and the TERT promoter. Probes targeting these genes were designed by Agilent Technologies. Furthermore, probes were also designed to target the HBV genome, which were used to detected HBV integrations in TERT gene (see Table S2 for detailed information). 10-200ng DNA samples were fragmented using Agilent's SureSelect Enzymatic Fragmentation Kit and subsequently used for library construction following the manufacturer's protocol. The targeted libraries were submitted for sequencing, achieving a mean depth of 1880X. Raw data processing involved the removal of low-quality reads and the trimming of adapters (60). The remained reads were mapped to the human genome (hg38) by BWA algorithm (61). SNVs were called by muTect (63) and varScan2(64). ANNOVAR (65) was employed to annotate mutations at the protein level to assess their potential functional effects. The protein-altering SNVs identified by either of the two tools were further manually checked.

Of note, 11 cancer-undetermined DNs in this study were subjected to both WES and panel sequencing. CNAs and protein-altering SNVs were identified by analyzing the WES data, TERT promoter alterations (promoter mutation and HBV integration) were further identified by analyzing the panel sequencing data.

**Mutation signature analysis**

Somatic SNVs were classified to 96 possible mutation types based on the sequence context of mutated base (6 types of substitution × 4 types of 5' base × 4 types of 3' base). The mutation matrix was reconstructed by the optimal linear combination of known mutational signatures that frequently occur in liver cancers (72). Only signatures whose contributions larger than 10% are shown in the plot, the remained signatures are called “other”. Associations between mutation signatures and exogenous and endogenous risk factors were obtained from COSMIC database.

**Neoantigen Predictions**

To examine neo-epitopes, 8-11 mer peptides around the mutated sites were extracted. HLA genotyping was predicted based on the RNAseq data for all major and minor HLA Class I alleles using OptiType (73). The bind affinity of these neo-epitopes to corresponding HLA-A, HLA-B and HLA-C alleles was estimated with NetMHCpan-4.1. Neoantigens with a relative percentile rank <= 0.5% were classified as strong binders, and neoantigens with a percentile <= 2% were classified as weak binders.

**Multiplex immunohistochemistry and multispectral image analysis**

3-5 µm-thick slides were cut from the FFPE blocks, deparaffinized, rehydrated through an ethanol gradient and unmasked with heating in citrate sodium solution (PH6). Then slides were stained using PANO 7-plex IHC kit (Cat# 004100100, Panovue) according to manufacturer’s instruction. HerPar (DAKO, IR624, 60 min incubation), CD4 (ZSGB-BIO, ZM0418, 30 min incubation), CD8 (CST, CST70306, 30 min incubation), PD1 (CST, CST43248, over-night incubation) and PDL1 (CST, CST13684, 30 min incubation) antibodies were applied, followed by horseradish peroxidase-conjugated secondary antibody incubation for 15 min and tyramide signal amplification for 10 min. The slides were microwave heat-treated after each tyramide signal amplification. Nuclei were stained with DAPI for 5 min after all the antigens above had been labelled. For CD4 and FOXP3 (CST, CST98377), MPO (abcam, ab208670), CD163(CST, CST93498) and CD11c (abcam, ab52632) co-staining, we used a TSA fluorescein evaluation kit (Perkin Elmer) following the manufacturer's instruction. The slides were scanned using the PerkinElmer Vectra 3 System with identical exposure time. By using spectral libraries that were previously built from images stained for each fluorophore (monoplex), the multispectral images obtained were unmixed through the inForm Advanced Image Analysis software.

The entire area of the slides was stained and scanned. We captured high-resolution images from NCa, DN and veHCC region of each case, respectively. For NCas, we captured images only from the parenchymal area but not contained the fibrotic bands. The scoring method consisted of several automated steps: tissue categorization, cell segmentation and cell phenotyping. Using the integrated inForm image analysis software, multispectral images that were representative of different samples were selected and used to train the inForm software for tissue categorization, cell segmentation and cell phenotyping. The settings learnt from the training on the representative images from different samples were saved within an algorithm, which enabled batch analysis of all the tissue slides.

**Quantification and statistical analysis**

No statistical methods were used to predetermine sample size. The experiments were not randomized. The investigators were not blinded to allocation during experiments and outcome assessment. $p < 0.05$ was taken to indicate statistical significance unless otherwise specified.

### Supplementary Table legends

Table S1. Summary of sample coding and clinicopathological Information

Table S2. Summary of sequencing data and the information of genes for target sequencing

Table S3. Summary of Cancer functional Genes (CFG) and CFG alterations in DNs and veHCCs

Table S4. Summary of protein-altering SNVs

Table S5. The list of gene signatures

### Supplementary Figure legends

**Fig. S1. Histological characterization and tissue sampling for sequencing**

(A-P) Whole-slide hematoxylin and eosin (H&E) images of nodule-in-nodule DNs with veHCCs inside. White dashed lines represent veHCC regions, while black dashed lines indicate DN regions. Blue circles denote multiple regions that were microdissected for whole genome sequencing (WGS) and RNA sequencing (RNA-seq). RNA-seq was not conducted for P6_veHCC, P9_veHCC_R2, and P17_veHCC due to insufficient material. DN, dysplastic nodule. veHCC, very early hepatocellular carcinoma.

**Fig. S2. Histopathological characterization of evolutionarily related DNs and veHCCs**

(A) Representative H&E staining of NCa, DN and veHCC regions of P8 (upper panel), and representative immunohistochemical staining of HSP70, GS and GPC3 of NCa, DN and veHCC regions of P8 (bottom panels). The NCa and DN region stained negatively for HSP70, GS and GPC3. The veHCC2 region stained positively for all the three markers. Scale bars, 100 μm.

(B) A summary of immunohistochemical staining of GPC3, HSP70, GS and histopathological characteristics of DNs and veHCCs.

**Fig. S3. SNVs in DNs and veHCCs**

(A) Heatmaps show the relationship of somatic SNVs between DNs and their paired veHCCs. SNVs are represented by horizontal color bars, and the color of each bar represents the variant allele frequency (VAF) of each SNV in DN or veHCC. Blue bars represent SNVs detected by WGS, and grey bars represent SNVs not detected. The average percentage of overlapped SNVs for each DN region of P6, P8 and each veHCC region of P9 were 92.6%, 91.1% and 97.6%, respectively. R1, R2 and R3 indicate region 1, region 2 and region 3.

(B) Estimated proportional contributions of each mutational signature to somatic SNVs in DNs and veHCCs. 10 known mutational signatures that frequently occur in HCCs were analyzed (Letouzé, E. et al., 2017). Mutational signatures with contributions exceeding 10% are presented, while the remaining signatures are categorized as "other".

(C) Follow-up magnetic resonance imaging of P3. White arrow represents the same lesion during the follow-up surveillance.

**Fig. S4. Comparison of SNV profiles in 2 HCC samples using freshly frozen and FFPE tissues**

Venn diagrams show the number of overlapped and private somatic single nucleotide variations (SNVs) between the freshly frozen (FF) and FFPE sample from the same HCC tissue.

**Fig. S5. Cancer-related genes induced by CNAs in cancer-prone and undetermined DNs**

(A, B) Violin plots show the number of pan-cancer oncogenes (A) and tumor suppressor genes (TSGs) (B) in cancer-prone DNs and cancer-undetermined DNs. Statistical significance is determined by unpaired student's t-test (two-tailed).

(C) Bar plot shows the percentage of cancer-prone DNs and cancer-undetermined DNs that acquired Chr8q (MYC) gain.

**Fig. S6. Genomic characterization of DNs and veHCCs**

(A) The number of protein-altering SNVs in evolutionarily unrelated DN and eHCC of each case. Statistical significance is determined by unpaired student's t-test (two-tailed).

(B) The fraction of the genome altered in evolutionarily unrelated DN and eHCC for each case is calculated by summing the lengths of CNAs for each chromosome, dividing by the total length of each chromosome, and then dividing by 22. CNA profiles of evolutionary unrelated DNs and eHCCs were inferred from WES data using CNVkit. Statistical significance is determined by unpaired student's t-test (two-tailed).

(C) The number of total SNVs detected by WGS in evolutionarily related DN and eHCC of each case. Statistical significance is determined by paired student's t-test (two-tailed).

(D) Heatmap shows the alteration landscape of cancer functional genes (CFGs) in evolutionarily related DNs and veHCCs.

(E) Heatmap shows the CNAs in evolutionarily related DNs and their matched veHCCs. BIC-Seq returned log2 ratios between the observed and expected number of reads in the segments. Segments with log2 ratio $>0.2$ or $<-0.2$ were regarded as somatic CNAs.

Red indicates copy number gain, while blue indicates copy number loss.

(F) The fraction of the genome altered in DN and veHCC for each case is calculated by summing the lengths of copy number alterations (CNAs) for each chromosome, dividing by the total length of each chromosome, and then dividing by 22. Significant difference is determined by paired student's t-test (two-tailed).

(G) Line plots show the number of pan-cancer oncogenes and tumor suppressor genes (TSGs) in DNs and paired veHCCs. Significant difference is determined by paired student's t-test (two-tailed).

(H, I) Gene set enrichment analysis of chromosome instability-related signature (H) and chromosome breakage-related signature (I) in evolutionarily related DNs and their paired veHCCs, as demonstrated by pre-ranked Gene Set Enrichment Analysis (GSEA). NES, normalized enrichment score in GSEA, FDR, false discovery rate. FDR < 0.25 indicates significance.

(J-L) Gene set enrichment analysis of DNA damage-related gene sets in evolutionarily related DNs and their paired veHCCs, as demonstrated by pre-ranked GSEA. FDR < 0.25 indicates significance.

(M, N) Linear plots of stepwise Chr1p loss during the malignant transition from DN to veHCC in P6 (M) and P9 (N).

(O) Heatmap shows the expression of a list of 2 tumor suppressor genes, located in the region of Chr1p (15-28Mb). RNA-seq data were not available for P6_veHCC.

**Fig. S7. Transcriptomic dynamics during the carcinogenesis from DNs to**

**veHCCs**

(A) Volcano plot shows differentially expressed genes in cancer-prone and cancer-undetermined DNs. Those genes that are significantly unregulated in cancer-prone and cancer-undetermined DNs are highlighted by orange and blue, respectively. (Fold change > 2, adjusted P_value < 0.05).

(B) Pathways enriched in cancer-prone DNs. 20 genes exhibited significant upregulation in cancer-prone DNs when compared to cancer-undetermined DNs (Fold change > 2, adjusted P_value < 0.05).

(C) A list of 20 small nucleolar RNAs (snoRNAs) that were significantly upregulated in cancer-undetermined DNs when compared to cancer-prone DNs (Fold change > 2, adjusted P_value < 0.05).

(D) Line plots illustrate three major modules of gene expression dynamics during the progression of carcinogenesis from NCas to veHCCs (left panel). The pathways enriched in each module are depicted in the right panel (FDR < 0.05). FDR, false discovery rate.

(E) Bar plot shows the expression of CDKN2A in NCas and DNs of these evolutionarily related cases. Significant difference is determined by unpaired student's t-test (two-tailed).

(F) Bar plot shows the density of Ki67+ cells in NCas, DNs and veHCCs of these evolutionarily related cases. Statistical significance is determined by unpaired student's t-test (two-tailed).

(G, H) Volcano plots show the changes in the activities of Reactome metabolism-related pathways between NCas and DNs (G), as well as between DNs and veHCCs (H). Pathways exhibiting significantly increased activities in NCas, DNs or veHCCs are highlighted by different colors. 80 metabolism-related gene sets were obtained from Reactome database. The activity of each metabolism-related pathway in each sample was assessed using GSVA. Statistical significance is determined by paired student's t-test (two-tailed). FDR < 0.05 indicates significance.

(I) Heatmap shows the activities of inflammation associated pathways in NCas and DNs of evolutionarily unrelated cases. The activity of each inflammation associated pathway in each sample is determined by GSVA.

**Fig. S8. Evolving immune response from DN to veHCC**

(A, B) Violin plots show the number of SNVs and predicted neoantigens in non-inflamed and inflamed veHCCs. Statistical significance is determined by unpaired student's t-test (two-tailed).

(C) Heatmap shows the expression of immune response-related markers, including chemokines, HLA class II molecules and immune cell markers in immune inflamed and non-inflamed veHCCs. Fold change > 2, adjust P_value < 0.05.

(D) Heatmaps show the abundance of immune cells and stromal cells in DNs and their matched non-inflamed veHCCs. The abundance of different immune cells and stromal cells are estimated through the deconvolution of gene expression using MCP-counter method. Statistical significance is determined by paired student's t-test (two-tailed). *P < 0.05.

(E) Line plots show the expression of CD4, CD8A and CD11c in DNs and paired immune-inflamed veHCCs. Significance is determined by paired student's t-test (two-tailed).

(F) Heatmaps show the abundance of immune cells and stromal cells in DNs and their matched inflamed veHCCs. The abundance of different immune cells and stromal cells are estimated through the deconvolution of gene expression using MCP-counter method. Statistical significance is determined by paired student's t-test (two-tailed). *P < 0.05, **P < 0.01.

(G) Schematic representation of bile acid synthesis pathways.

(H) Gene set enrichment analysis of bile acid metabolism-related gene set in inflamed and non-inflamed veHCCs, as demonstrated by pre-ranked GSEA.

(I) Heatmaps show the abundance of immune cells and stromal cells in NCas and their paired inflamed veHCCs. Statistical significance is determined by paired student's t-test (two-tailed). *P < 0.05, **P < 0.01.

(J) Line plots show the densities of CD4+ T cells and CD8+ T cells in NCas and their paired inflamed veHCCs. The densities of CD4+ and CD8+ T cells are determined by multiplex staining of CD4 and CD8. Statistical significance is determined by paired student's t-test (two-tailed).

(K) Gene set enrichment analysis of angiogenesis-associated TAM signature and M2 macrophage signature in DNs and related inflamed veHCCs, as demonstrated by pre-ranked GSEA. TAM, tumor associated macrophage.

(L) Dot plot shows the GSVA scores of TGF-beta signature in DNs and their matched inflamed veHCCs. Significance is determined by paired student's t-test (two-tailed).

(M) Line plot shows the densities of CD163+ macrophages in DNs and their paired inflamed veHCCs. Statistical significance is determined by paired student's t-test (two-tailed).

**Fig. S9. Phylogenetic trees of tumorigenesis**

Schematic illustration of the progression of carcinogenesis from NCas to veHCCs. Key events in the genesis of cancer-prone DNs including TERT alterations (82%), Chr7p/7q gain (24%) and inactive immune response. 2 major evolutionary scenarios of the malignant transition from DNs to veHCCs were illustrated, the accumulation of high burden of CNAs (Scenario1) and an immune-inflamed status with immune evasion (Scenario 2).

**Fig. S10. Phylogenetic trees of tumorigenesis**

Phylogenetic trees of P11, P13, P14, P16 and P19 (A, Scenario 1), P5, P9, P10, P17 and P20 (B, Scenario 2), P7 and P8 (C, Scenario 1 and Scenario 2). Point mutations or CNAs of CFGs were manually annotated on the trees. The trunk and branch lengths are proportional to the number of SNVs.

**Fig. S11. Two evolutionary scenarios of the malignant transition of P7_DN**

(A) Schematic illustration of the 2 evolutionary scenarios of the malignant transition from P7_DN1 to P7_veHCC1 and P7_veHCC2 (Scenario 1), as well as malignant transition from P7_DN2 to P7_veHCC3 (Scenario 2). HBV integration in TERT gene and point mutations or CNAs of other CFGs were manually annotated on the trees.

(B) Bar plot shows the CNA burden (fraction of genome altered) in the DN and paired veHCCs of P7.

(C) Representative H&E staining (upper panel), multiplex staining of HepPar, PD1, PDL1, CD4, CD8 (middle panel) and multiplex staining of CD163, MPO, CD11c (bottom panel) of P7. Scale bars, 100 μm.

(D) Bar plots show the densities of CD4+ and CD8+ T cells, CD11c+ DC cells, CD163+ macrophage, MPO+ neutrophil, PD1+ CD8+ T cells, PDL1+ cells and FOXP3+ CD4+ T cells in DNs and veHCCs of P7.

### Figure S1

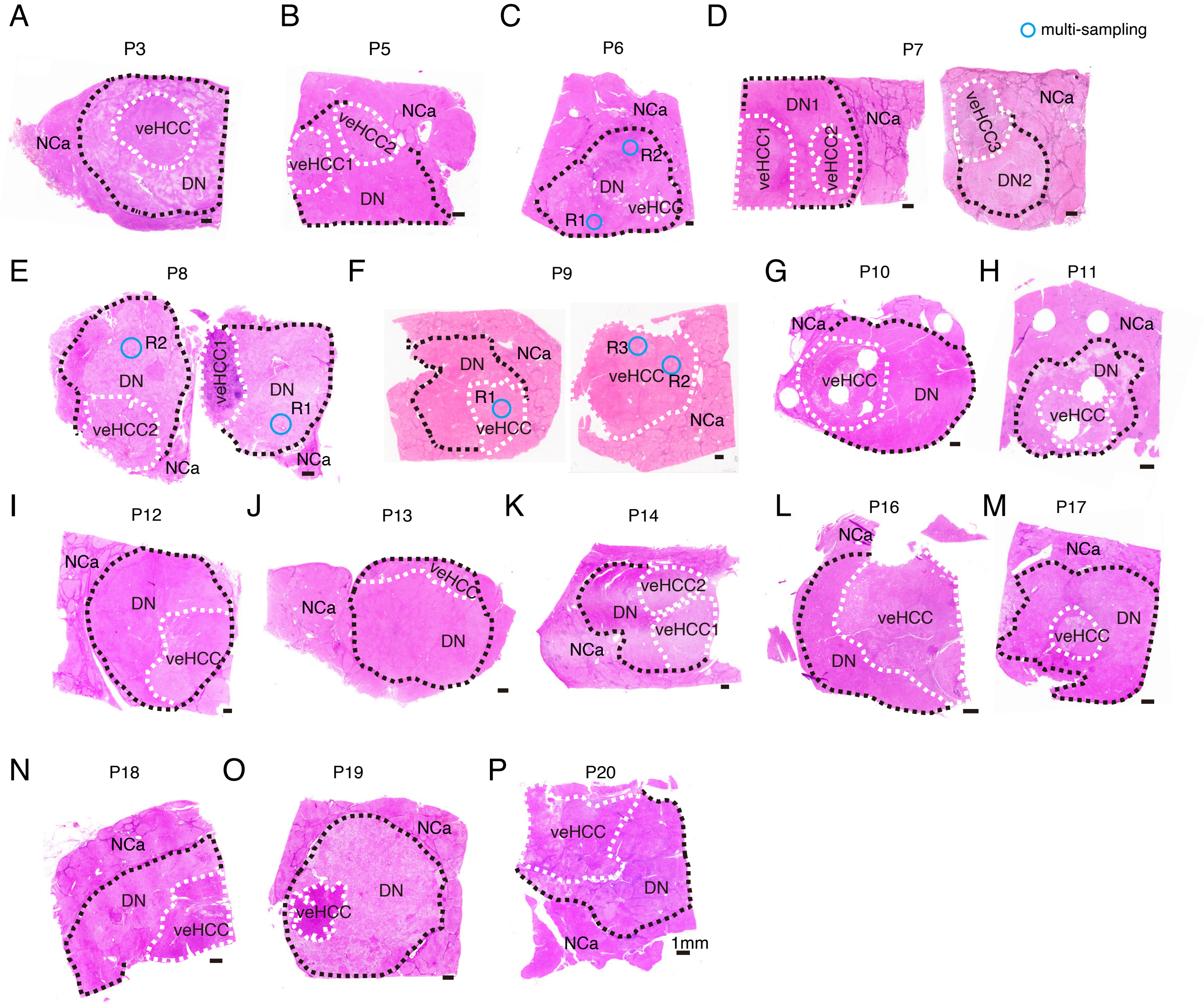
A
P3
NCa
veHCC
DN
B
P5
NCa
veHCC2
veHCC1
DN
C
P6
NCa
R2
DN
veHCC
R1
D
P7
multi-sampling
DN1
NCa
veHCC1
veHCC2
veHCC3
NCa
DN2
E
P8
R2
DN
veHCC2
NCa
veHCC1
DN
R1
NCa
F
P9
DN
NCa
R1
veHCC
R3
veHCC
R2
NCa
G
P10
NCa
veHCC
DN
H
P11
NCa
DN
veHCC
I
P12
NCa
DN
veHCC
J
P13
veHCC
NCa
DN
K
P14
veHCC2
DN
veHCC1
NCa
L
P16
NCa
veHCC
DN
M
P17
NCa
DN
veHCC
N
P18
NCa
DN
veHCC
O
P19
NCa
DN
veHCC
P
P20
veHCC
DN
NCa
1mm

### Figure S2

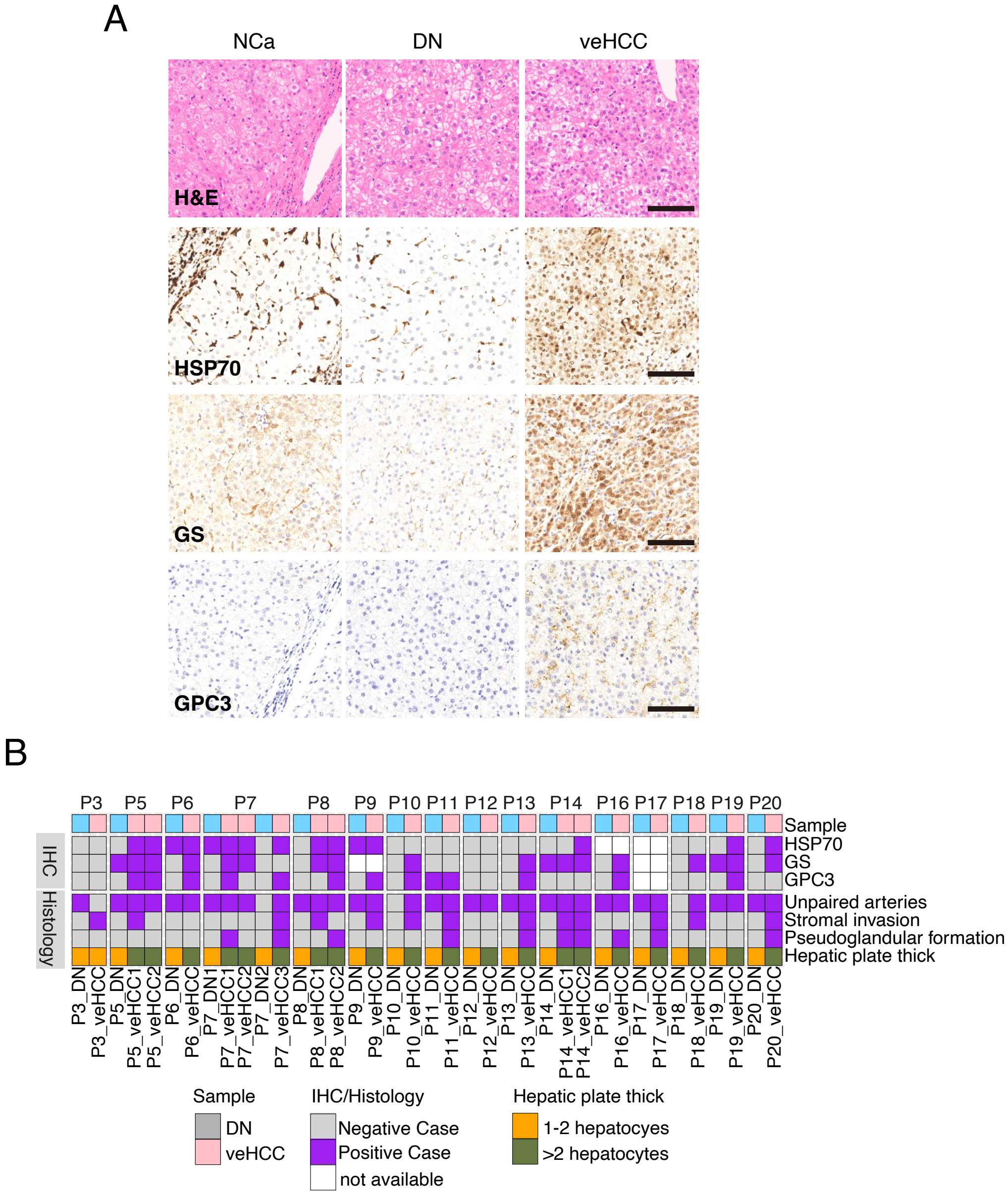
A
NCa
DN
veHCC
H&E
HSP70
GS
GPC3
B
P3
P5
P6
P7
P8
P9
P10
P11
P12
P13
P14
P16
P17
P18
P19
P20
Sample
HSP70
GS
GPC3
Unpaired arteries
Stromal invasion
Pseudoglandular formation
Hepatic plate thick
IHC
Histology
P3_DN
P3_veHCC
P5_DN
P5_veHCC1
P5_veHCC2
P6_DN
P6_veHCC
P7_DN1
P7_veHCC1
P7_veHCC2
P7_DN2
P7_veHCC3
P8_DN
P8_veHCC1
P8_veHCC2
P9_DN
P9_veHCC
P10_DN
P10_veHCC
P11_DN
P11_veHCC
P12_DN
P12_veHCC
P13_DN
P13_veHCC
P14_DN
P14_veHCC1
P14_veHCC2
P16_DN
P16_veHCC
P17_DN
P17_veHCC
P18_DN
P18_veHCC
P19_DN
P19_veHCC
P20_DN
P20_veHCC
Sample
DN
veHCC
IHC/Histology
Negative Case
Positive Case
not available
Hepatic plate thick
1-2 hepatocytes
>2 hepatocytes

### Figure S3

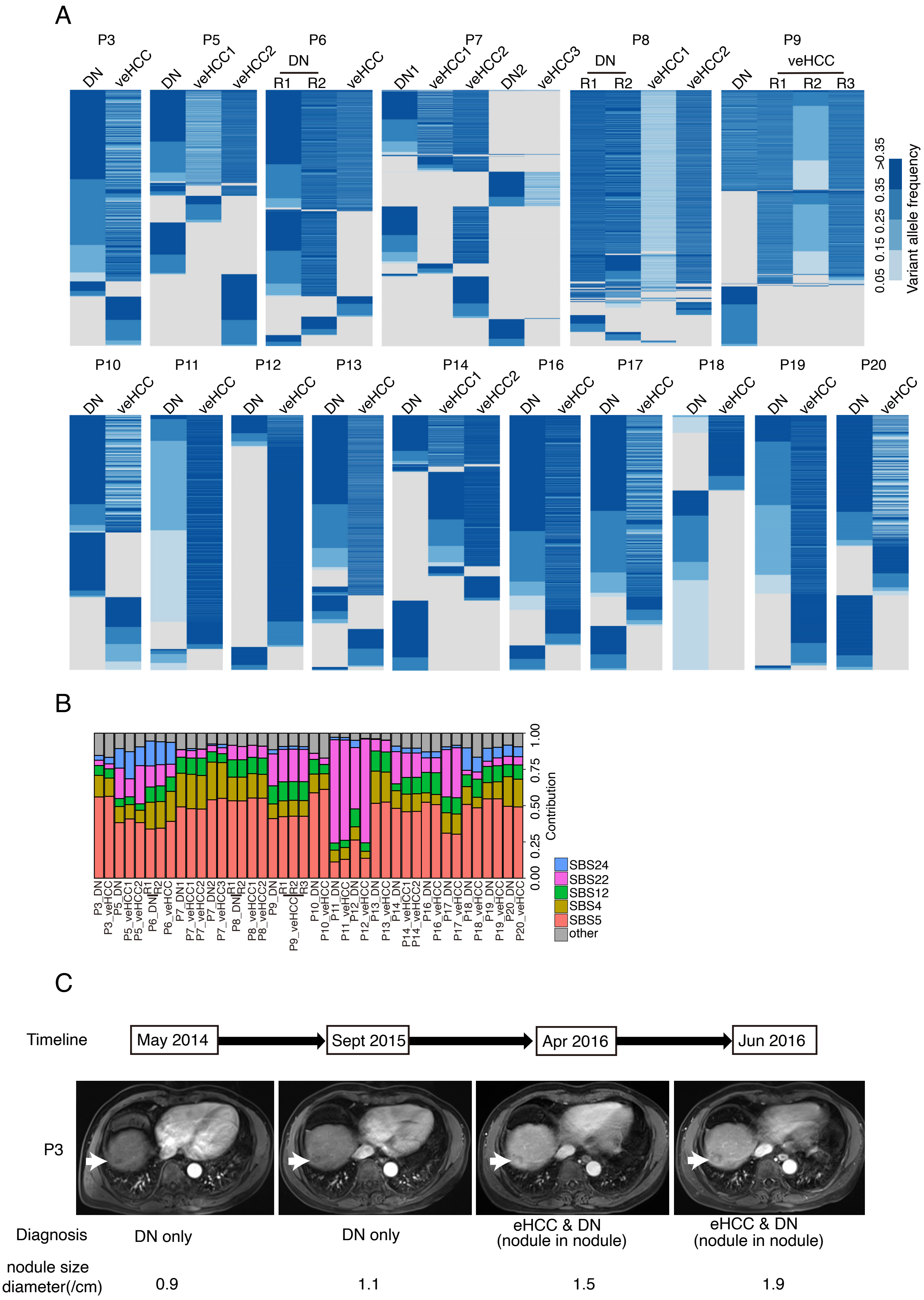
A
P3
DN
veHCC
P5
DN
veHCC1
veHCC2
P6
DN
R1
R2
veHCC
P7
DN1
veHCC1
veHCC2
DN2
veHCC3
P8
DN
R1
R2
veHCC1
veHCC2
P9
DN
veHCC
R1
R2
R3
0.05 0.15 0.25 0.35 >0.35
Variant allele frequency
P10
P11
P12
P13
P14
P16
P17
P18
P19
P20
B
Contribution
0.00 0.25 0.50 0.75 1.00
SBS24
SBS22
SBS12
SBS4
SBS5
other
C
Timeline
May 2014
Sept 2015
Apr 2016
Jun 2016
P3
Diagnosis
DN only
DN only
eHCC & DN
(nodule in nodule)
eHCC & DN
(nodule in nodule)
nodule size
diameter(/cm)
0.9
1.1
1.5
1.9

### Figure S4

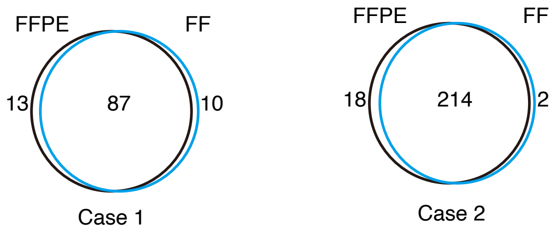

### Figure S5

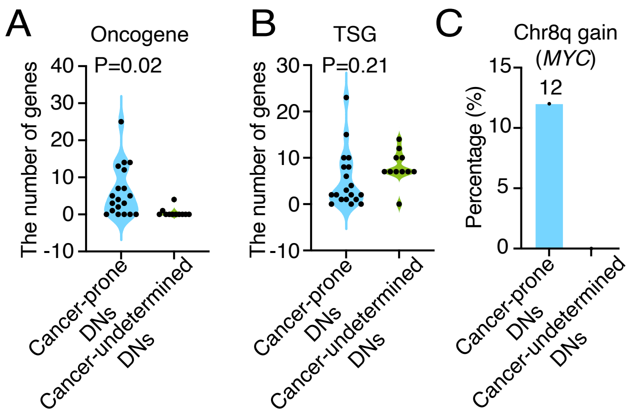
A
Oncogene
P=0.02
The number of genes
40
30
20
10
0
-10
Cancer-prone DNs
Cancer-undetermined DNs
B
TSG
P=0.21
The number of genes
30
20
10
0
-10
Cancer-prone DNs
Cancer-undetermined DNs
C
Chr8q gain (*MYC*)
12
Percentage (%)
15
10
5
0
Cancer-prone DNs
Cancer-undetermined DNs

### Figure S6

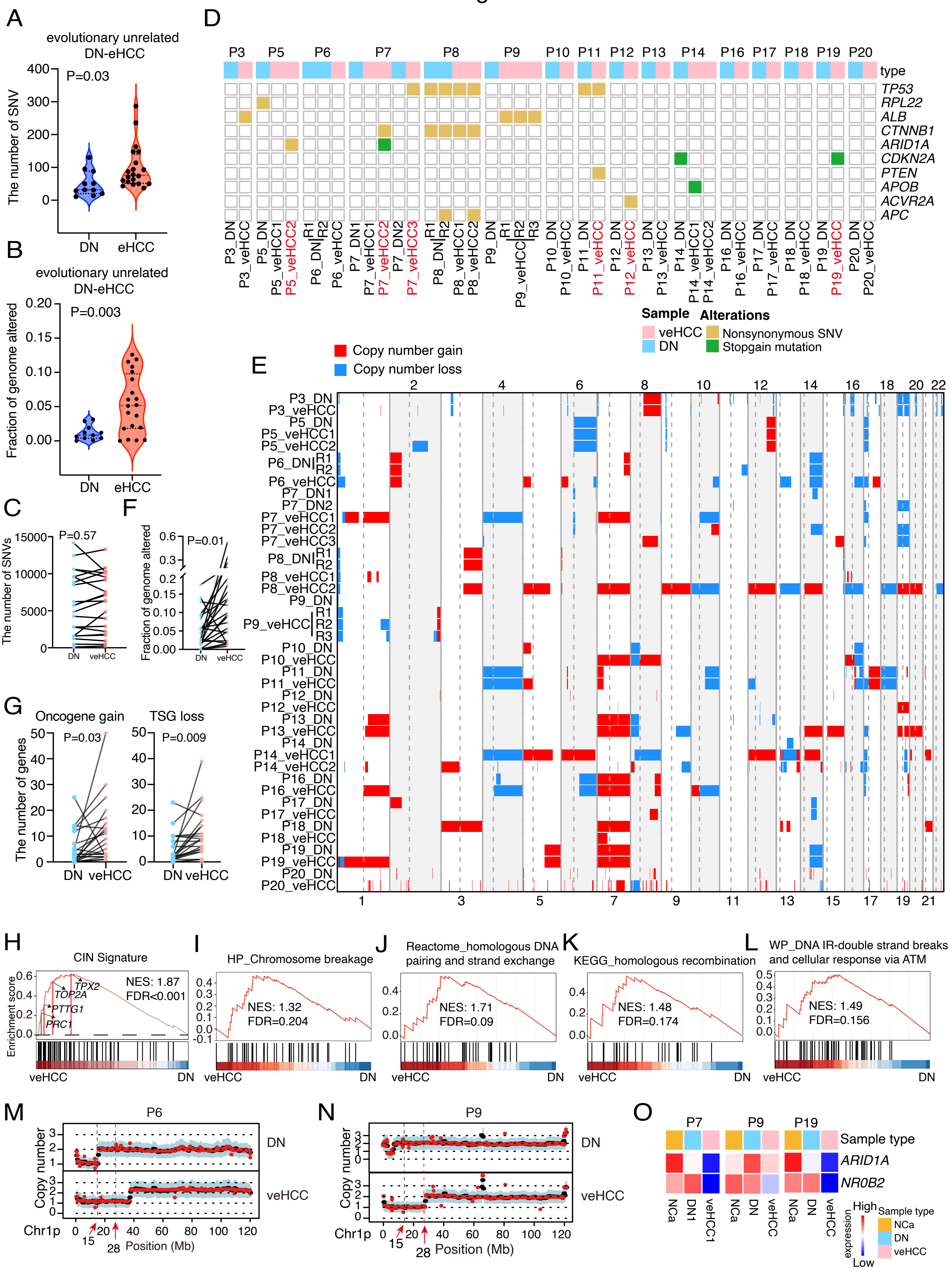

A
evolutionary unrelated DN-eHCC
P=0.03
The number of SNV
DN
eHCC
B
evolutionary unrelated DN-eHCC
P=0.003
Fraction of genome altered
C
P=0.57
The number of SNVs
F
P=0.01
Fraction of genome altered
G
Oncogene gain
P=0.03
TSG loss
P=0.009
The number of genes
DN veHCC
D
type
TP53
RPL22
ALB
CTNNB1
ARID1A
CDKN2A
PTEN
APOB
ACVR2A
APC
Sample
veHCC
DN
Alterations
Nonsynonymous SNV
Stopgain mutation
E
Copy number gain
Copy number loss
H
CIN Signature
NES: 1.87
FDR<0.001
TPX2
TOP2A
PTTG1
PRC1
Enrichment score
I
HP_Chromosome breakage
NES: 1.32
FDR=0.204
J
Reactome_homologous DNA pairing and strand exchange
NES: 1.71
FDR=0.09
K
KEGG_homologous recombination
NES: 1.48
FDR=0.174
L
WP_DNA IR-double strand breaks and cellular response via ATM
NES: 1.49
FDR=0.156
veHCC
DN
M
P6
N
P9
Copy number
Chr1p
Position (Mb)
O
P7
P9
P19
Sample type
ARID1A
NR0B2
NCa
DN
veHCC
High
Low
expression

### Figure S7

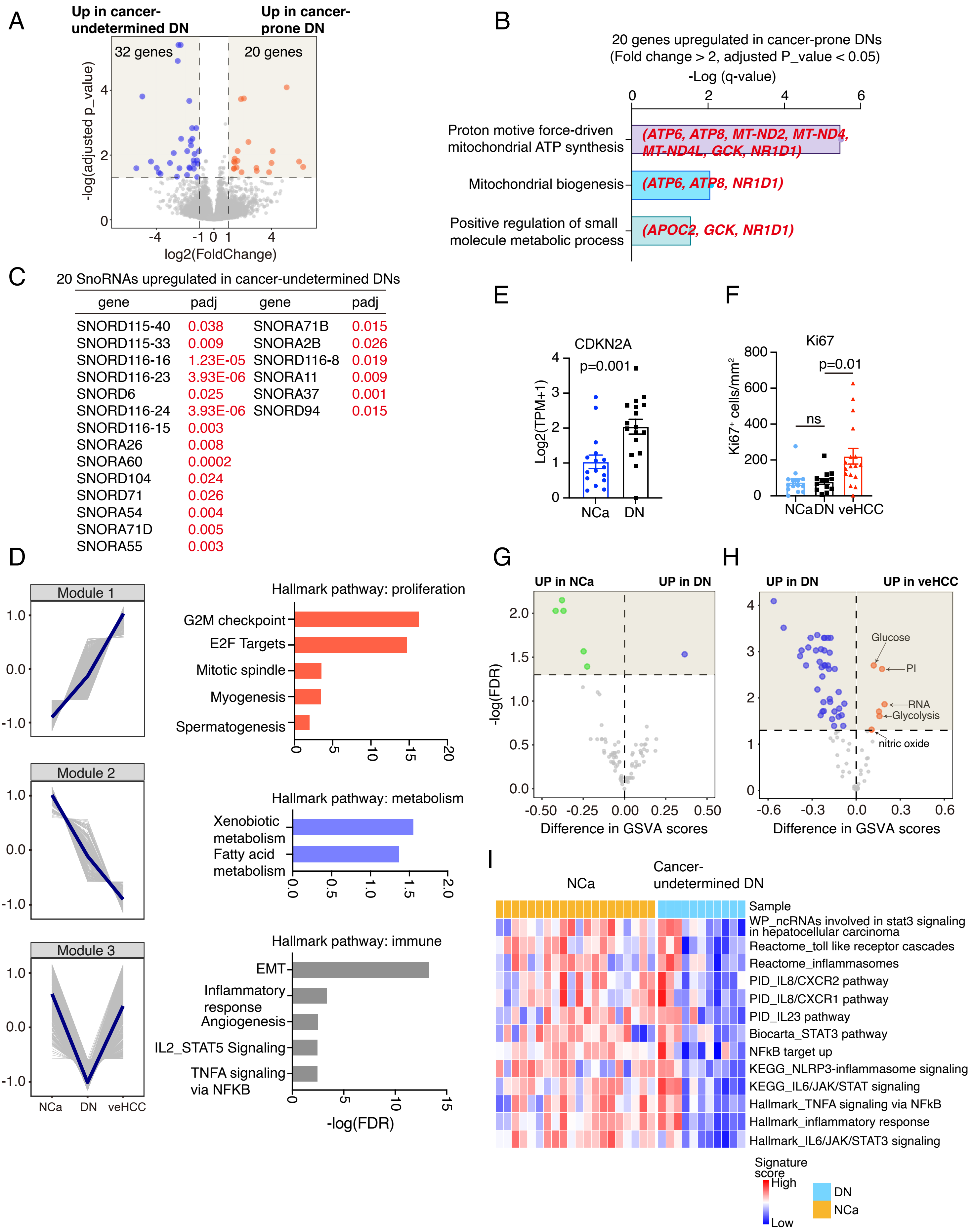


| gene | padj | gene | padj |
|---|---|---|---|
| SNORD115-40 | 0.038 | SNORA71B | 0.015 |
| SNORD115-33 | 0.009 | SNORA2B | 0.026 |
| SNORD116-16 | 1.23E-05 | SNORD116-8 | 0.019 |
| SNORD116-23 | 3.93E-06 | SNORA11 | 0.009 |
| SNORD6 | 0.025 | SNORA37 | 0.001 |
| SNORD116-24 | 3.93E-06 | SNORD94 | 0.015 |
| SNORD116-15 | 0.003 | | |
| SNORA26 | 0.008 | | |
| SNORA60 | 0.0002 | | |
| SNORD104 | 0.024 | | |
| SNORD71 | 0.026 | | |
| SNORA54 | 0.004 | | |
| SNORA71D | 0.005 | | |
| SNORA55 | 0.003 | | |

### Figure S8

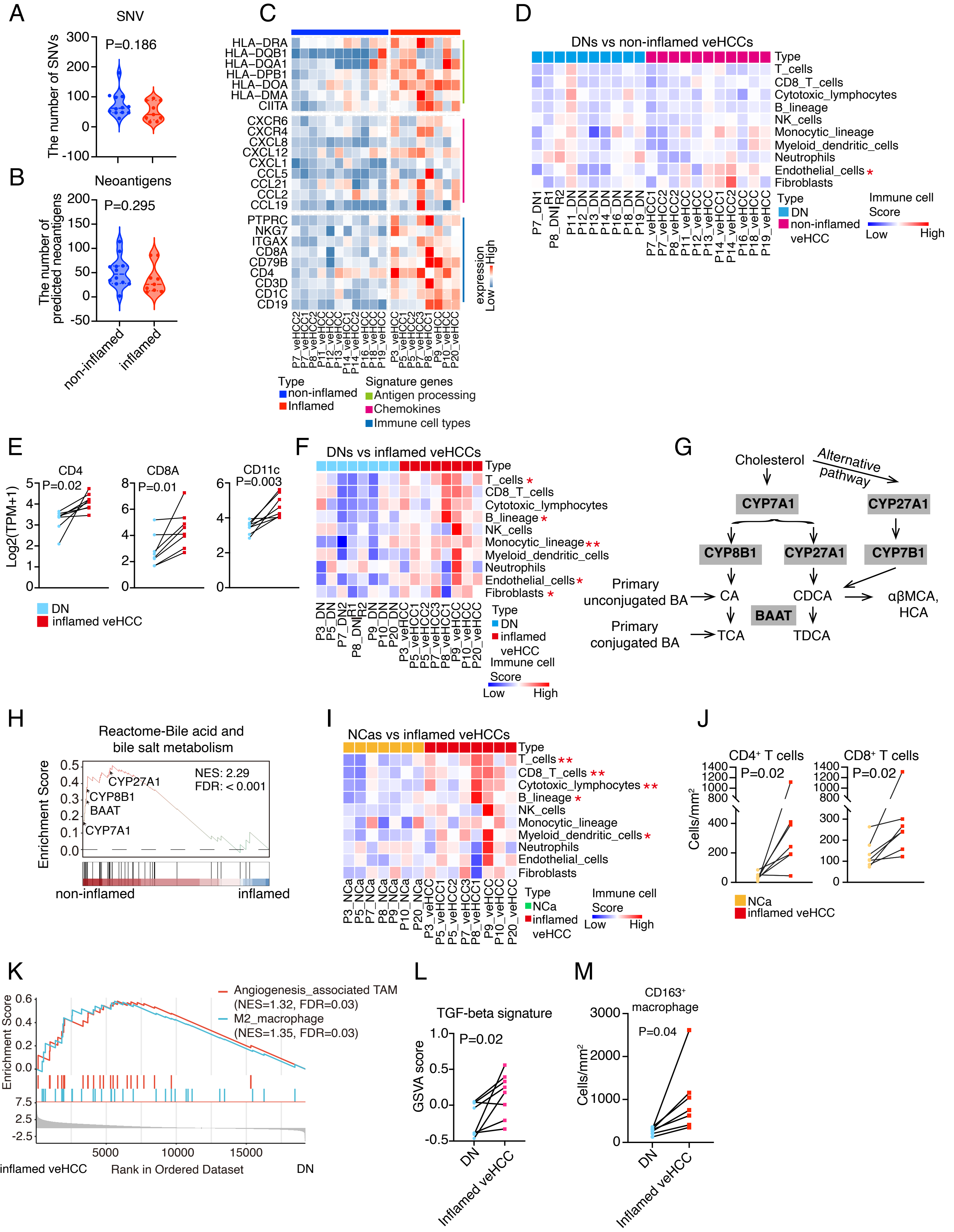
A
SNV
P=0.186
The number of SNVs
B
Neoantigens
P=0.295
The number of predicted neoantigens
non-inflamed
inflamed
C
HLA−DRA
HLA−DQB1
HLA−DQA1
HLA−DPB1
HLA−DOA
HLA−DMA
CIITA
CXCR6
CXCR4
CXCL8
CXCL12
CXCL1
CCL5
CCL21
CCL2
CCL19
PTPRC
NKG7
ITGAX
CD8A
CD79B
CD4
CD3D
CD1C
CD19
expression
Low
High
Type
non-inflamed
Inflamed
Signature genes
Antigen processing
Chemokines
Immune cell types
D
DNs vs non-inflamed veHCCs
Type
T_cells
CD8_T_cells
Cytotoxic_lymphocytes
B_lineage
NK_cells
Monocytic_lineage
Myeloid_dendritic_cells
Neutrophils
Endothelial_cells *
Fibroblasts
Type
DN
non-inflamed veHCC
Immune cell Score
Low
High
E
CD4
P=0.02
CD8A
P=0.01
CD11c
P=0.003
Log2(TPM+1)
DN
inflamed veHCC
F
DNs vs inflamed veHCCs
Type
T_cells *
CD8_T_cells
Cytotoxic_lymphocytes
B_lineage *
NK_cells
Monocytic_lineage **
Myeloid_dendritic_cells
Neutrophils
Endothelial_cells *
Fibroblasts *
Type
DN
inflamed veHCC
Immune cell Score
Low
High
G
Cholesterol
Alternative pathway
CYP7A1
CYP27A1
CYP8B1
CYP27A1
CYP7B1
Primary unconjugated BA
CA
CDCA
αβMCA, HCA
BAAT
Primary conjugated BA
TCA
TDCA
H
Reactome-Bile acid and bile salt metabolism
NES: 2.29
FDR: < 0.001
CYP27A1
CYP8B1
BAAT
CYP7A1
Enrichment Score
non-inflamed
inflamed
I
NCas vs inflamed veHCCs
Type
T_cells **
CD8_T_cells **
Cytotoxic_lymphocytes **
B_lineage *
NK_cells
Monocytic_lineage
Myeloid_dendritic_cells *
Neutrophils
Endothelial_cells
Fibroblasts
Type
NCa
inflamed veHCC
Immune cell Score
Low
High
J
CD4⁺ T cells
P=0.02
CD8⁺ T cells
P=0.02
Cells/mm²
NCa
inflamed veHCC
K
Angiogenesis_associated TAM (NES=1.32, FDR=0.03)
M2_macrophage (NES=1.35, FDR=0.03)
Enrichment Score
inflamed veHCC
Rank in Ordered Dataset
DN
L
TGF-beta signature
P=0.02
GSVA score
DN
Inflamed veHCC
M
CD163⁺ macrophage
P=0.04
Cells/mm²
DN
Inflamed veHCC

### Figure S9

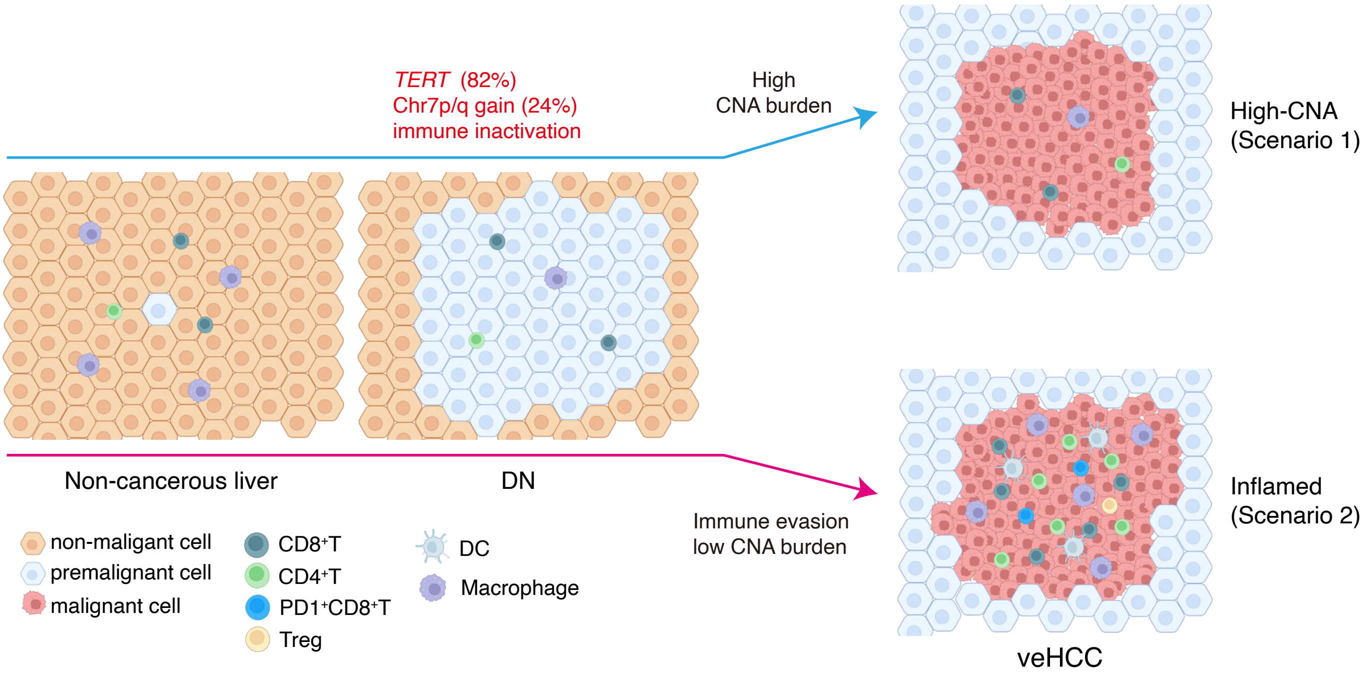

*TERT* (82%)
Chr7p/q gain (24%)
immune inactivation
High
CNA burden
High-CNA
(Scenario 1)
Non-cancerous liver
DN
Immune evasion
low CNA burden
Inflamed
(Scenario 2)
veHCC
non-maligant cell
premalignant cell
malignant cell
CD8⁺T
CD4⁺T
PD1⁺CD8⁺T
Treg
DC
Macrophage

### Figure S10

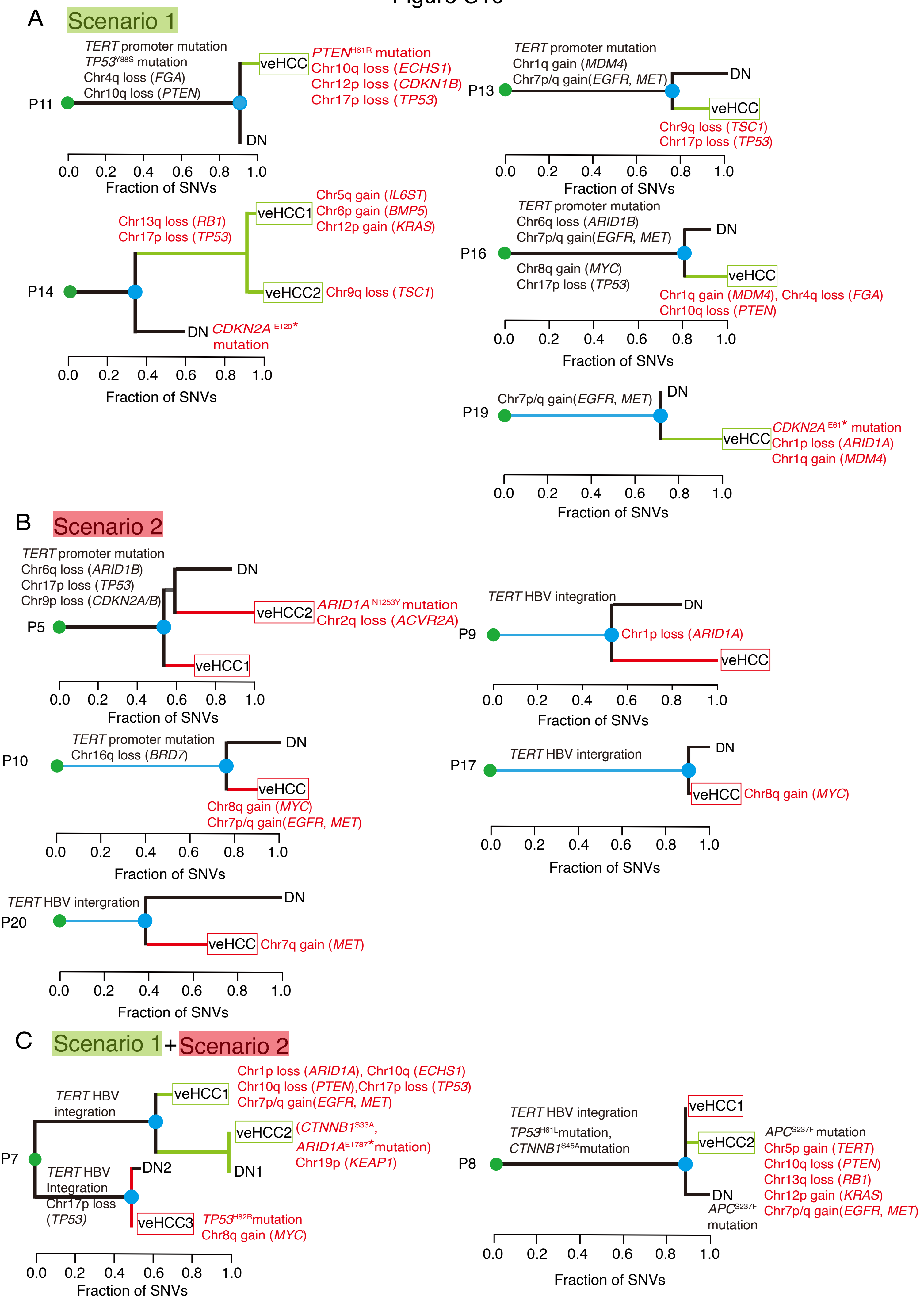

A
Scenario 1
P11
TERT promoter mutation
TP53 Y88S mutation
Chr4q loss (FGA)
Chr10q loss (PTEN)
veHCC
PTEN H61R mutation
Chr10q loss (ECHS1)
Chr12p loss (CDKN1B)
Chr17p loss (TP53)
DN
0.0 0.2 0.4 0.6 0.8 1.0
Fraction of SNVs
P13
TERT promoter mutation
Chr1q gain (MDM4)
Chr7p/q gain(EGFR, MET)
DN
veHCC
Chr9q loss (TSC1)
Chr17p loss (TP53)
0.0 0.2 0.4 0.6 0.8 1.0
Fraction of SNVs
P14
Chr13q loss (RB1)
Chr17p loss (TP53)
veHCC1
Chr5q gain (IL6ST)
Chr6p gain (BMP5)
Chr12p gain (KRAS)
veHCC2
Chr9q loss (TSC1)
DN
CDKN2A E120* mutation
0.0 0.2 0.4 0.6 0.8 1.0
Fraction of SNVs
P16
TERT promoter mutation
Chr6q loss (ARID1B)
Chr7p/q gain(EGFR, MET)
Chr8q gain (MYC)
Chr17p loss (TP53)
DN
veHCC
Chr1q gain (MDM4), Chr4q loss (FGA)
Chr10q loss (PTEN)
0.0 0.2 0.4 0.6 0.8 1.0
Fraction of SNVs
P19
Chr7p/q gain(EGFR, MET)
DN
veHCC
CDKN2A E61* mutation
Chr1p loss (ARID1A)
Chr1q gain (MDM4)
0.0 0.2 0.4 0.6 0.8 1.0
Fraction of SNVs
B
Scenario 2
P5
TERT promoter mutation
Chr6q loss (ARID1B)
Chr17p loss (TP53)
Chr9p loss (CDKN2A/B)
DN
veHCC2
ARID1A N1253Y mutation
Chr2q loss (ACVR2A)
veHCC1
0.0 0.2 0.4 0.6 0.8 1.0
Fraction of SNVs
P9
TERT HBV integration
DN
Chr1p loss (ARID1A)
veHCC
0.0 0.2 0.4 0.6 0.8 1.0
Fraction of SNVs
P10
TERT promoter mutation
Chr16q loss (BRD7)
DN
veHCC
Chr8q gain (MYC)
Chr7p/q gain(EGFR, MET)
0.0 0.2 0.4 0.6 0.8 1.0
Fraction of SNVs
P17
TERT HBV intergration
DN
veHCC
Chr8q gain (MYC)
0.0 0.2 0.4 0.6 0.8 1.0
Fraction of SNVs
P20
TERT HBV intergration
DN
veHCC
Chr7q gain (MET)
0.0 0.2 0.4 0.6 0.8 1.0
Fraction of SNVs
C
Scenario 1 + Scenario 2
P7
TERT HBV integration
veHCC1
Chr1p loss (ARID1A), Chr10q (ECHS1)
Chr10q loss (PTEN),Chr17p loss (TP53)
Chr7p/q gain(EGFR, MET)
veHCC2
(CTNNB1 S33A, ARID1A E1787* mutation)
Chr19p (KEAP1)
DN1
TERT HBV Integration
Chr17p loss (TP53)
DN2
veHCC3
TP53 H82R mutation
Chr8q gain (MYC)
0.0 0.2 0.4 0.6 0.8 1.0
Fraction of SNVs
P8
TERT HBV integration
TP53 H61L mutation,
CTNNB1 S45A mutation
veHCC1
veHCC2
APC S237F mutation
Chr5p gain (TERT)
Chr10q loss (PTEN)
Chr13q loss (RB1)
Chr12p gain (KRAS)
Chr7p/q gain(EGFR, MET)
DN
APC S237F mutation
0.0 0.2 0.4 0.6 0.8 1.0
Fraction of SNVs

### Figure S11

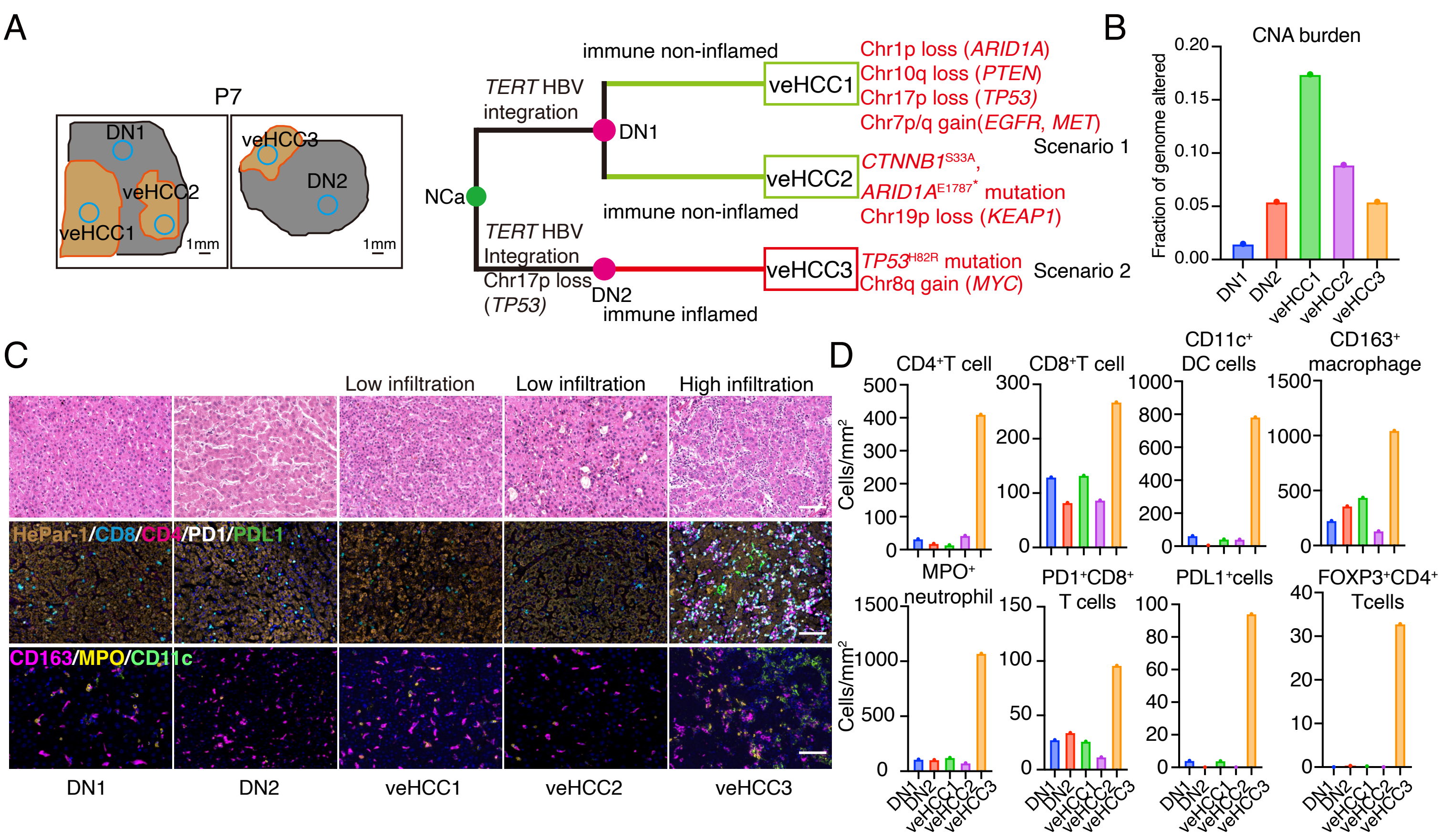
A
P7
DN1
veHCC3
veHCC2
DN2
veHCC1
1mm
1mm
immune non-inflamed
TERT HBV integration
DN1
veHCC1
veHCC2
NCa
immune non-inflamed
TERT HBV Integration
Chr17p loss (TP53)
DN2
veHCC3
immune inflamed
Chr1p loss (ARID1A)
Chr10q loss (PTEN)
Chr17p loss (TP53)
Chr7p/q gain(EGFR, MET)
Scenario 1
CTNNB1 S33A,
ARID1A E1787* mutation
Chr19p loss (KEAP1)
TP53 H82R mutation
Chr8q gain (MYC)
Scenario 2
B
CNA burden
Fraction of genome altered
0.20
0.15
0.10
0.05
0.00
DN1
DN2
veHCC1
veHCC2
veHCC3
C
Low infiltration
Low infiltration
High infiltration
HePar-1/CD8/CD4/PD1/PDL1
CD163/MPO/CD11c
DN1
DN2
veHCC1
veHCC2
veHCC3
D
CD4+T cell
CD8+T cell
CD11c+ DC cells
CD163+ macrophage
Cells/mm2
MPO+ neutrophil
PD1+CD8+ T cells
PDL1+cells
FOXP3+CD4+ Tcells
DN1
DN2
veHCC1
veHCC2
veHCC3
